# Genetic analysis of mitochondrial DNA copy number and associated traits identifies loci implicated in nucleotide metabolism, platelet activation, and megakaryocyte proliferation, and reveals a causal association of mitochondrial function with mortality

**DOI:** 10.1101/2021.01.25.428086

**Authors:** RJ Longchamps, SY Yang, CA Castellani, W Shi, J Lane, ML Grove, TM Bartz, C Sarnowski, K Burrows, AL Guyatt, TR Gaunt, T Kacprowski, J Yang, PL De Jager, L Yu, CHARGE Aging and Longevity Group, A Bergman, R Xia, M Fornage, MF Feitosa, MK Wojczynski, AT Kraja, MA Province, N Amin, F Rivadeneira, H Tiemeier, AG Uitterlinden, L Broer, JBJ Van Meurs, CM Van Duijn, LM Raffield, L Lange, SS Rich, RN Lemaitre, MO Goodarzi, CM Sitlani, ACY Mak, DA Bennett, S Rodriguez, JM Murabito, KL Lunetta, N Sotoodehnia, G Atzmon, Y Kenny, N Barzilai, JA Brody, BM Psaty, KD Taylor, JI Rotter, E Boerwinkle, N Pankratz, DE Arking

## Abstract

Mitochondrial DNA copy number (mtDNA-CN) measured from blood specimens is a minimally invasive marker of mitochondrial function that exhibits both inter-individual and intercellular variation. To identify genes involved in regulating mitochondrial function, we performed a genome-wide association study (GWAS) in 465,809 White individuals from the Cohorts for Heart and Aging Research in Genomic Epidemiology (CHARGE) consortium and the UK Biobank (UKB). We identified 133 SNPs with statistically significant, independent effects associated with mtDNA-CN across 100 loci. A combination of fine-mapping, variant annotation, and co-localization analyses were used to prioritize genes within each of the 133 independent sites. Putative causal genes were enriched for known mitochondrial DNA depletion syndromes (*p* = 3.09 x 10^−15^) and the gene ontology (GO) terms for mtDNA metabolism (*p* = 1.43 x 10^−8^) and mtDNA replication (*p* = 1.2 x 10^−7^). A clustering approach leveraged pleiotropy between mtDNA-CN associated SNPs and 41 mtDNA-CN associated phenotypes to identify functional domains, revealing three distinct groups, including platelet activation, megakaryocyte proliferation, and mtDNA metabolism. Finally, using mitochondrial SNPs, we establish causal relationships between mitochondrial function and a variety of blood cell related traits, kidney function, liver function and overall (*p* = 0.044) and non-cancer mortality (*p* = 6.56 x 10^−4^).

## Introduction

Mitochondria are the cellular organelles primarily responsible for producing the chemical energy required for metabolism, as well as signaling the apoptotic process, maintaining homeostasis, and synthesizing several macromolecules such as lipids, heme and iron-sulfur clusters^1,2^. Mitochondria possess their own genome (mtDNA); a circular, intron-free, double-stranded, haploid, ~16.6 kb maternally inherited molecule encoding 37 genes vital for proper mitochondrial function. Due to the integral role of mitochondria in cellular metabolism, mitochondrial dysfunction is known to play a critical role in the underlying etiology of several aging-related diseases^3–5^.

Unlike the nuclear genome, a large amount of variation exists in the number of copies of mtDNA present within cells, tissues, and individuals. The relative copy number of mtDNA (mtDNA-CN) has been shown to be positively correlated with oxidative stress^6^, energy reserves, and mitochondrial membrane potential^7^. As a minimally invasive proxy measure of mitochondrial dysfunction^8^, decreased mtDNA-CN measured in blood has been previously associated with aging-related disease states including frailty^9^, cardiovascular disease^10–12^, chronic kidney disease^13^, neurodegeneration^14,15^, and cancer^16^.

Although mtDNA-CN measured from whole blood presents itself as an easily accessible and minimally invasive biomarker, cell type composition has been shown to be an important confounder, complicating analyses^17,18^. For example, while platelets generally have fewer mtDNA molecules than leukocytes, the lack of a platelet nuclear genome drastically skews mtDNA-CN estimates. As a result, not only is controlling for cell composition extremely vital for accurate mtDNA-CN estimation, interpreting the results in relation to the impact of cell composition becomes a necessity^18–20^.

Although the comprehensive mechanism through which mtDNA-CN is modulated is largely unknown^21,22^, twin studies have estimated a broad-sense heritability of ~0.65, consistent with moderate genetic control^23^. Several nuclear genes have been shown to directly modulate mtDNA-CN, specifically those within the mtDNA replication machinery such as the mitochondrial polymerase, *POLG* and *POLG2*^24,25^, as well as the mitochondrial DNA helicase, *TWNK*, and the mitochondrial single-stranded binding protein, *mtSSB*^26^. Furthermore, nuclear genes which maintain proper mitochondrial nucleotide supply including *DGUOK* and *TK2* have also been shown to regulate mtDNA-CN^27–29^. To further elucidate the genetic control over mtDNA-CN, several genome-wide association studies (GWAS) of mtDNA-CN have been published^30–33^, including a study that was published while the current manuscript was in preparation, analyzing ~300,000 participants from the UK Biobank (UKB), and identifying 50 independent loci^33^.

In the present study, we report mtDNA-CN GWAS results from 465,809 individuals across the Cohorts for Heart and Aging Research in Genomic Epidemiology (CHARGE) consortium^34^ and the UK Biobank (UKB)^35^. Using multiple gene prioritization and functional annotation methods, we assign genes to loci that reach genome-wide significance. We perform a PHEWAS and group our genome-wide significant SNPs into 3 clusters that represent distinct functional domains related to mtDNA-CN. Finally, we leverage mitochondrial SNPs to establish causality between mitochondrial function and mtDNA-CN associated traits.

## Subjects and Methods

### Study Populations

470,579 individuals participated in this GWAS, 465,809 of whom self-identified as White. Participants were derived from 7 population-based cohorts representing the Cohorts for Heart and Aging Research in Genetic Epidemiology (CHARGE) consortium (Avon Longitudinal Study of Parents and Children [ALSPAC], Atherosclerosis Risk in Communities [ARIC], Cardiovascular Health Study [CHS], Multi-Ethnic Study of Atherosclerosis [MESA], Religious Orders Study and Memory and Aging Project [ROSMAP], Study of Health in Pomerania [SHIP]) and from the UK Biobank (UKB) (Supplemental Table 1). Detailed descriptions of each participating cohort, their quality control practices, study level analyses, and ethic statements are available in the Supplemental Methods. All study participants provided written informed consent and all centers obtained approval from their institutional review boards.

### Methods for Mitochondrial DNA Copy Number Estimation (CHARGE cohorts)

#### qPCR

mtDNA-CN was determined using a quantitative PCR assay as previously described^32,36^. Briefly, the cycle threshold (Ct) value of a nuclear-specific and mitochondrial-specific probe were measured in triplicate for each sample. In CHS, a multiplex assay using the mitochondrial *ND1* probe and nuclear *RPPH1* probe was used, whereas ALSPAC used a mitochondrial probe targeting the D-Loop and a nuclear probe targeting *B2M*. In CHS, we observed plate effects, as well as a linear increase in ΔCt due to the pipetting order of each replicate. These effects were corrected in the analysis using linear mixed model regression, with pipetting order included as a fixed effect and plate as a random effect to create a raw measure of mtDNA-CN. In ALSPAC, run-to-run variability was controlled using 3 calibrator samples added to every plate, to allow for adjustment by a per-plate calibration factor^32^.

#### Microarray

Microarray probe intensities were used to estimate mtDNA-CN using the Genvisis software package^37^ as previously described^10,36^. Briefly, Genvisis uses the median mitochondrial probe intensity across all homozygous mitochondrial SNPs as an initial estimate of mtDNA-CN. Technical artifacts such as DNA input quality, DNA input quantity, and hybridization efficiency were captured through either surrogate variable (SV) or principal component (PC) analyses. SVs or PCs were adjusted for through stepwise linear regression by adding successive components until each successive surrogate variable or principal component no longer significantly improved the model.

#### Whole Genome Sequencing (ARIC)

Whole genome sequencing read counts were used to estimate mtDNA-CN as previously described^36^. Briefly, the total number of reads in a sample were web scraped from the NCBI sequence read archive. Mitochondrial reads were downloaded directly from dbGaP through Samtools (1.3.1). There was no overlap between ARIC microarray and ARIC whole-genome sequencing samples. A ratio of mitochondrial reads to total aligned reads was used as a raw measure of mtDNA-CN.

#### Adjusting for Covariates

Each method described above represents a raw measure of mtDNA-CN, adjusted for technical artifacts; however, several potential confounding variables (e.g., age, sex, blood cell composition) have been identified previously^18^. Raw mtDNA-CN values were adjusted for white blood cell count in ARIC, SHIP and CHS (which also adjusted for platelet count), depending on available data. Standardized residuals (mean = 0, standard deviation = 1) of mtDNA-CN were used after adjusting for covariates (Supplemental Table 1).

### Estimation of Mitochondrial DNA Copy Number (UKB)

Due to the availability of more detailed cell count data, as well as a different underlying biochemistry for the Affymetrix Axiom array compared to the genotyping arrays used in the CHARGE cohorts, mtDNA-CN in the UKB was estimated differently (Supplemental Methods). Briefly, mtDNA-CN estimates derived from whole exome sequencing data, available on ~50,000 individuals, were generated first using customized Perl scripts to aggregate the number of mapped sequencing reads and correct for covariates through both linear and spline regression models. Concurrently, mitochondrial probe intensities from the Affymetrix Axiom arrays, available on the full ~500,000 UKB cohort, were adjusted for technical artifacts through principal components generated from nuclear probe intensities. Probe intensities were then regressed onto the whole exome sequencing mtDNA-CN metric, and beta estimates from that regression were used to estimate mtDNA-CN in the full UKB cohort. Finally, we used a 10-fold cross validation method to select the cell counts to include in the final model (Supplemental Table 2). The final UKB mtDNA-CN metric is the standardized residuals (mean = 0, standard deviation = 1) from a linear model adjusting for covariates (age, sex, cell counts) as described in the Supplemental Methods.

### Genome-Wide Association Study

For each individual cohort, regression analysis was performed with residualized mtDNA-CN as the dependent variable adjusting for age, sex, and cohort-specific covariates (e.g., principal components, DNA collection site, family structure, cell composition). Cohorts with multiple mtDNA-CN estimation platforms were stratified into separate analyses. Ancestry-stratified meta-analyses were performed using Metasoft software using the Han and Eskin random effects model to control for unobserved heterogeneity due to differences in mtDNA-CN estimation method^38^. Effect size estimates for SNPs were calculated using a random effect meta-analysis from cohort summary statistics, as the Han and Eskin model relaxes the assumption under the null hypothesis without modifying the effect size estimates that occur under the alternative hypothesis^38^. In total, three complementary analyses were performed in self-identified White individuals: (1) a meta-analysis using all available studies, (2) a meta-analysis of studies with available data for cell count adjustments, and (3) an analysis of UKB-only data. As the vast majority of samples are derived from the UKB study, and given the difficulty in interpreting effect size estimates from a random effects model, further downstream analyses were all performed using effect size estimates from UKB-only data. We additionally performed X chromosome analyses, using only UKB data. X-chromosome analyses were stratified by sex (males = 194,151, females = 216,989), and summary statistics were meta-analyzed using METAL^39^ to obtain the final effect estimates.

### SNP Heritability Estimation

SNP heritability estimates were retrieved from BOLT-LMM^40^. To verify this metric, we used SumHer^41^ to calculate an independent heritability metric using summary statistics. The heritability model used in this analysis was the BLD-LDAK model. The tagging file used is the pre-computed UK Biobank GBR version for the corresponding heritability model. The summary statistics were filtered so that only single-character reference and alternate alleles are allowed. Chr:BP combination duplicates were removed except the first appearance. SNP heritability was then calculated and extracted from output files.

### Identification of Independent GWAS Loci

To identify the initial genome-wide significant (lead) SNPs in each locus, the most significant SNP that passed genome-wide significance (*p* < 5 x 10^−8^) within a 1 Mb window was selected. To avoid Type I error, SNPs were only retained for further analyses if there were either (a) at least two genome-wide significant SNPs in the 1 Mb window or (b) if the lead SNP was directly genotyped. Conditional analyses were performed in UKB, where the lead SNPs from the original GWAS were used as additional covariates in order to identify additional independent associations.

### Comparisons with Hägg et al. 2020

To compare results with Hägg et al. 2020^33^, summary statistics were obtained from their Supplementary Table 4. Loci were identified as shared between the two GWAS if two lead SNPs were fewer than 500,000 base pairs apart from one another.

### Fine-mapping

The susieR package was used to identify all potential causal variants for each independent locus associated with mtDNA CN^42^. UKB imputed genotype data for unrelated White subjects were used and variants were extracted using a 500 kb window around the lead SNP for each locus with minor allele frequency (MAF) > 0.001. 95% credible sets (CS) of SNPs, containing a potential causal variant within a locus, were generated. The minimum absolute correlation within each CS is 0.5 and the scaled prior variance is 0.01. When the CS did not include the lead SNP identified from the GWAS, some of the parameters were slightly relaxed [minimum absolute correlation is 0.2, estimate prior variance is TRUE]. The SNP with the highest posterior inclusion probability (PIP) within each CS was also identified (Supplemental Table 3). With a few exceptions, final lead SNPs were selected by prioritizing initially identified SNPs unless the SNP with the highest PIP had a PIP greater than 0.2 and was 1.75 times larger than the SNP with the second highest PIP.

### Functional Annotation and Gene Prioritization

#### Functional Annotation

ANNOVAR was used for functional annotation of variants identified in the fine-mapping step^43^. First, variants were converted to an ANNOVAR-ready format using the dbSNP version 150 database^44^. Then, variants were annotated with ANNOVAR using the RefSeq Gene database^45^. The annotation for each variant includes the associated gene and region (e.g., exonic, intronic, intergenic). For intergenic variants, ANNOVAR provides flanking genes and the distance to each gene. For exonic variants, annotations also include likely functional consequences (e.g., synonymous/nonsynonymous, insertion/deletion), the gene affected by the variant, and the amino acid sequence change (Supplemental Table 4).

#### Co-localization Analyses

Co-localization analyses were performed using the approximate Bayes factor method in the R package *coloc*^46^. Briefly, *coloc* utilizes eQTL data and GWAS summary statistics to evaluate the probability that gene expression and GWAS data share a single causal SNP (colocalize). *Coloc* returns multiple posterior probabilities; H0 (no causal variant), H1 (causal variant for gene expression only), H2 (causal variant for mtDNA-CN only), H3 (two distinct causal variants), and H4 (shared causal variant for gene expression and mtDNA-CN). In the event of high H4, we designate the gene as causal for the GWAS phenotype of interest (mtDNA-CN). eQTL summary statistics were obtained from the eQTLGen database^47^. Genes with significant associations with lead SNPs were tested for co-localization using variants within a 500 kb window of the sentinel SNP. Occasionally, some of the eQTLGen p-values for certain SNPs were identical due to R’s (ver 4.0.3) limitation in handling small numbers. To account for this, if the absolute value for a SNP’s z-score association with a gene was greater than 37.02, z-scores were rescaled so that the largest z-score would result in a p-value of 5 x 10^−300^. Additionally, a few clearly co-localized genes did not result in high H4 PPs due to the strong effect for each phenotype of a single SNP (Supplemental Figure 1), possibly due to differences in linkage disequilibrium (LD) between the UKB and eQTLGen populations. To account for this, we summed mtDNA-CN GWAS p-values and eQTLGen p-values for each SNP and removed the SNP with the lowest combined p-value. Co-localization analyses were then repeated without the lowest SNP. Genes with H4 greater than 50% were classified as genes with significant evidence of co-localization.

#### DEPICT

Gene prioritization was performed with Depict, an integrative tool that incorporates gene co-regulation and GWAS data to identify the most likely causal gene at a given locus^48^. Across GWAS SNPs which overlapped with the DEPICT database, we identified SNPs representing 119 independent loci with LD pruning defined as *p* < 5 x 10^−8^, r^2^ < 0.05 and > 500 kb from other locus boundaries. Only genes with a nominal p-value of less than 0.05 were considered for downstream prioritization.

#### Gene Assignment

To prioritize genes for each identified locus, we utilized functional annotations, eQTL co-localization analyses, and DEPICT gene prioritization results (Supplemental Figure 2). First, genes with missense variants within susieR fine-mapped credible sets were assigned to loci. If loci co-localized with a gene’s expression with a posterior probability (PP) of greater than 0.50 and there were no other co-localized genes with a PP within 5%, the gene with the highest posterior probability was assigned. If there was still no assigned gene, the most significant DEPICT gene was assigned. If there was no co-localization or DEPICT evidence, the nearest gene was assigned.

### Gene Set Enrichment Analyses

Using the finalized gene list from the prioritization pipeline, GO and KEGG pathway enrichment analyses were performed using the “goana” and “kegga” functions from the R package *limma*^49^, treating all known genes as the background universe^50^. Only one gene per locus was used for “goana” and “kegga” gene set enrichment analysis, prioritizing genes assigned to primary independent hits. If there were multiple assigned genes, one gene was randomly selected to avoid biasing results through loci with multiple functionally related genes. To identify an appropriate p-value cutoff, 100 genes were randomly selected from the genome and run through the same enrichment analysis. This permutation was repeated 1000 times to generate a null distribution of the smallest p-values from each permutation. For cluster-specific gene set enrichment analyses, permutation testing used the same number of random genes as the number of genes in each cluster. To ensure robustness of results, gene set enrichment analysis was repeated 50 times with random selection of genes at loci with multiple assigned genes. GO and KEGG terms that passed permutation cutoffs at least 40/50 times were retained.

### Gene-based Association Test

We used metaXcan, which employs gene expression prediction models to evaluate associations between phenotypes and gene expression^51^. We obtained pre-calculated expression prediction models and SNP covariance matrices, computed using whole blood from European ancestry individuals in version 7 of the Genotype-Tissue expression (GTEx) database^52^. Using prediction performance *p* < 0.05, a total of 6,285 genes were predicted. Of these genes, 74 passed Bonferroni correction of *p* < 7.95 x 10^−6^. Gene set enrichment analyses were performed on Bonferroni-significant genes as previously described. REVIGO^53^ was used on the “medium” setting (allowed similarity = 0.7) to visualize significantly enriched GO terms.

We used a one-sided Fisher’s exact test to test for enrichment of genes that have been previously identified as causal for mtDNA depletion syndromes^54–56^.

### PHEWAS-based SNP Clustering

#### mtDNA-CN Phenome-wide Association Study (PHEWAS)

We used the PHEnome Scan ANalysis Tool (PHESANT)^57^ to identify mtDNA-CN associated quantitative traits in the UKB. Briefly, we tested for the association of mtDNA-CN with 869 quantitative traits (Supplemental Table 5), limiting analyses to 365,781 White, unrelated individuals (used.in.pca.calculation=1). As extreme cell count measurements could indicate individuals with active infections or cancers, they were excluded from analysis (see Supplemental Methods). Analyses were adjusted for age, sex, and assessment center.

#### SNP-Phenotype Associations

SNP genotypes were regressed on mtDNA-associated quantitative phenotypic traits using linear regression, adjusted for sex, age with a natural spline (df = 2), assessment center, genotyping array, and 40 genotyping principal components (provided as part of the UKB data download).

#### SNP Clustering

To identify distinct clusters of mtDNA-CN GWS SNPs based on phenotypic associations, beta estimates from the SNP-phenotype associations were first divided by the beta estimate of the mtDNA-CN SNP-mtDNA-CN association, so that all SNP-phenotype associations are relative to the mtDNA-CN increasing allele and scaled to the effect of the SNP on mtDNA-CN. The adjusted beta estimates were subjected to a dimensionality reduction method, Uniform Manifold and Approximation Projection (UMAP), as implemented in the R package *umap*^58^ (random_state=123, n_neighbors=10, min_dist=0.001, n_components=2, n_epochs=200). SNPs were assigned to clusters using Density Based Clustering of Applications with Noise (DBSCAN) as implemented in the R package *dbscan*^59^ (minPts=10). Robustness of cluster assignment was established by varying n_neighors, min_dist, and random_state parameters. Clusters represent groups of SNPs with similar phenotypic associations.

#### Phenotype Enrichment and Permutation Testing

To test for enrichment of *individual* phenotypes within clusters, we compared the median mtDNA-CN scaled phenotype beta estimates within the cluster to the median beta estimates for all SNPs not in the cluster, with significance determined using 20,000 permutations in which cluster assignment was permuted. For multi-test correction *across all* phenotypes, we performed 300 permutations of the initial cluster assignment, followed by the comparison of median beta estimates as described above. We retained only the most significant result from across all phenotypes and clusters from each of the 300 permutations, and then selected the 15^th^ most significant value as the study-wide threshold for multi-test corrected significance of p < 0.05.

### mtDNA Variant Association Analyses

#### Mitochondrial Variant Phasing and Imputation

Shapeit4 and Impute5 were used for UK Biobank mtDNA genotype phasing and imputation^60,61^. Phasing and imputation were performed separately for each genotyping array (UKBB, UKBL), and restricted to self-identified White individuals. The reference panel used for imputation analysis was the 1000 Genomes Project phase 3 mtDNA variants^62^. UK Biobank genotypes were coded to match the reference panel allele. All genotype files, including the reference panel, were phased using Shapeit4 to fill in any missing genotypes using the phasing iteration sequence “10b,1p,1b,1p,1b,1p,1b,1p,10m”, where b is burn-in iteration, p is pruning iteration, and m is main iteration. The ‒sequencing option was also used due to the presence of multiple mtDNA variants in a very small region, analogous to sequencing data.

Phased UK Biobank genotypes were then imputed with the reference panel using Impute5 with the following parameters: ‒pbwt-depth 8; ‒pbwt-cm 0.005; ‒no-threshold. All imputed variants were functionally annotated using MSeqDR mvTools^63^.

### mtSNP Association Tests

Linear regressions stratified by genotyping array (UKBB, UKBL) were performed for each mtDNA SNP on the 41 traits and mtDNA-CN, including the following covariates: age, age^2^, sex, center, first 20 genotyping PCs. Only SNPs with MAF > 0.005 and imputation INFO score > 0.80 were included (UKBB, n=223; UKBL, n=190; both, n=149). Results were then meta-analyzed using inverse variance weighting. For association analyses between mitochondrial SNPs/haplotypes and mtDNA-CN, the mtDNA-CN metric used was derived only from WES data, as mtDNA genotypes can subtly influence the estimates from genotype array intensities.

### Identification of Independent Genetic Effects

Single SNP study-wide significance was established by generating 300 normally distributed dummy traits, and running single SNP tests using the UKBB data. The minimum SNP p-value for each dummy trait was then selected, and the 15^th^ most significant p-value from the 300 analyses was divided by 42 (41 real traits + mtDNA-CN), resulting in study-wide p-value threshold of *p* < 9.5×10^−6^. To identify a subset of traits to perform credible set identification using SusieR (see above, **Fine Mapping**), SNPs were first filtered based on the study-wide p-value threshold, and then then most significantly associated trait was identified for each SNP. SusieR, (parameters: L=10, estimate_residual_variance=TRUE, estimate_prior_variance=TRUE, check_z = FALSE) was then run for each of these traits using the UKBB imputed data and summary association test statistics. A total of 7 credible sets were identified across the 4 traits, two of which co-localized, resulting in 6 credible sets. Independence across the 6 credible sets was tested using multivariate regression models, and requiring *p* < 0.0005 for at least one trait for a SNP to remain in the model. SNPs MT73A_G and MT 7028C_T were in moderate high LD (r^2^=0.67), but based on conditional regression analyses as described in the main results, capture independent effects and are associated with different traits.

### Haplotype Generation and Analysis

Haplotype were constructed by concatenating SNPs across the 6 credible sets using SNPs directly genotyped on both genotyping arrays. This required selecting a SNP with a lower PIP for 2 of the 6 credible sets (MT12612A_G replaced MT462C_T, r^2^ = 0.81; MT10238T_C replaced MT4529A_T, r^2^ = 0.89). Haplotypes with MAF < 0.005 were set to missing (n = 1607), resulting in 8 haplotypes, with the most common haplotype set as reference. Significance for haplotype associations with each trait were generated by an anova between regression models with and without the haplotypes. Covariates included age, age^2^, sex, center, first 40 genotyping PCs, and genotyping array.

Mortality analyses were run using Cox proportional hazards models, with covariates as above. Individuals with external causes of death (ICD 10 Death Code categories V, W, X, Y) were censored at time of death. Additionally, for non-cancer mortality analyses, cancer death (ICD 10 Death Code categories C00-D48) were censored. For cancer mortality analyses, all death due to non-cancer cases were censored at time of death.

Clustering for visualization was performed using the R package ‘heatmaply’, with default setting and hclust_method=”ward.D2”.

All statistical analyses were performed using R version 4.0.3.

## Results

### Sample Characteristics

The current study included 465,809 White individuals (53.9% female) with an average age of 56.6 yrs (sd = 8.2 yrs) (Supplemental Table 1). Follow-up validation analyses were performed in 4,770 Blacks (60.2% female) with an average age of 61.2 yrs (sd = 7.4 yrs). The majority of the data originated from the UKB (93%). The bulk of the DNA used for mtDNA-CN estimation was derived from buffy coat (95.5%) while the rest was derived from peripheral leukocytes (2.2%), whole blood (2.3%), or brain (< 0.2%). mtDNA-CN estimated from Affymetrix genotyping arrays consisted of 97.9% of the data while the remainder was derived from qPCR (1.8%) and WGS (0.3%).

### GWAS

Previous work has demonstrated that the method used to measure mtDNA-CN can impact the strength of association^36^. To account for potential differences across studies due to the different mtDNA-CN measurements used, as well as the inclusion of blood cell counts as covariates in only a subset of the cohorts, we took two approaches. First, we used a random effects model to perform meta-analyses, allowing for different genetic effect size estimates across cohorts. Second, we performed three complementary analyses in individuals who self-identified as White: 1) meta-analysis of all available studies (n = 465,809); 2) meta-analysis of studies with available data for cell count adjustment (n = 456,151); and 3) GWAS of UKB only (n = 440,266) (Figure 1). 77 loci were significant in all three meta-analyses, and we identified 93 independent loci that were significant in at least one of the analyses. In the meta-analysis of UKB-only data, 92 of the total 93 loci were identified (Supplemental Figure 3). Given that >90% of the samples come from the UKB study, and the challenge of interpreting effect size estimates from a random effects model, downstream analyses all use effect size estimates from the UKB only analyses (Supplemental Table 6), which showed no evidence for population substructure inflating test statistics, with a genomic inflation factor of 1.09 (Supplemental Figure 4). SNP heritability estimated from BOLT-LMM^40^ for mtDNA-CN adjusted for age and sex was 10.5% while heritability for mtDNA-CN adjusted for age, sex, and cell counts was 7.4%, implying that some of the mtDNA-CN heritability observed in previous studies could be due to heritability of cell type composition. We also used SumHer^41^ as an alternative approach to calculating SNP heritability, which returned a comparable estimate of 7.0% for the cell-count corrected mtDNA-CN metric.

**Figure 1.**
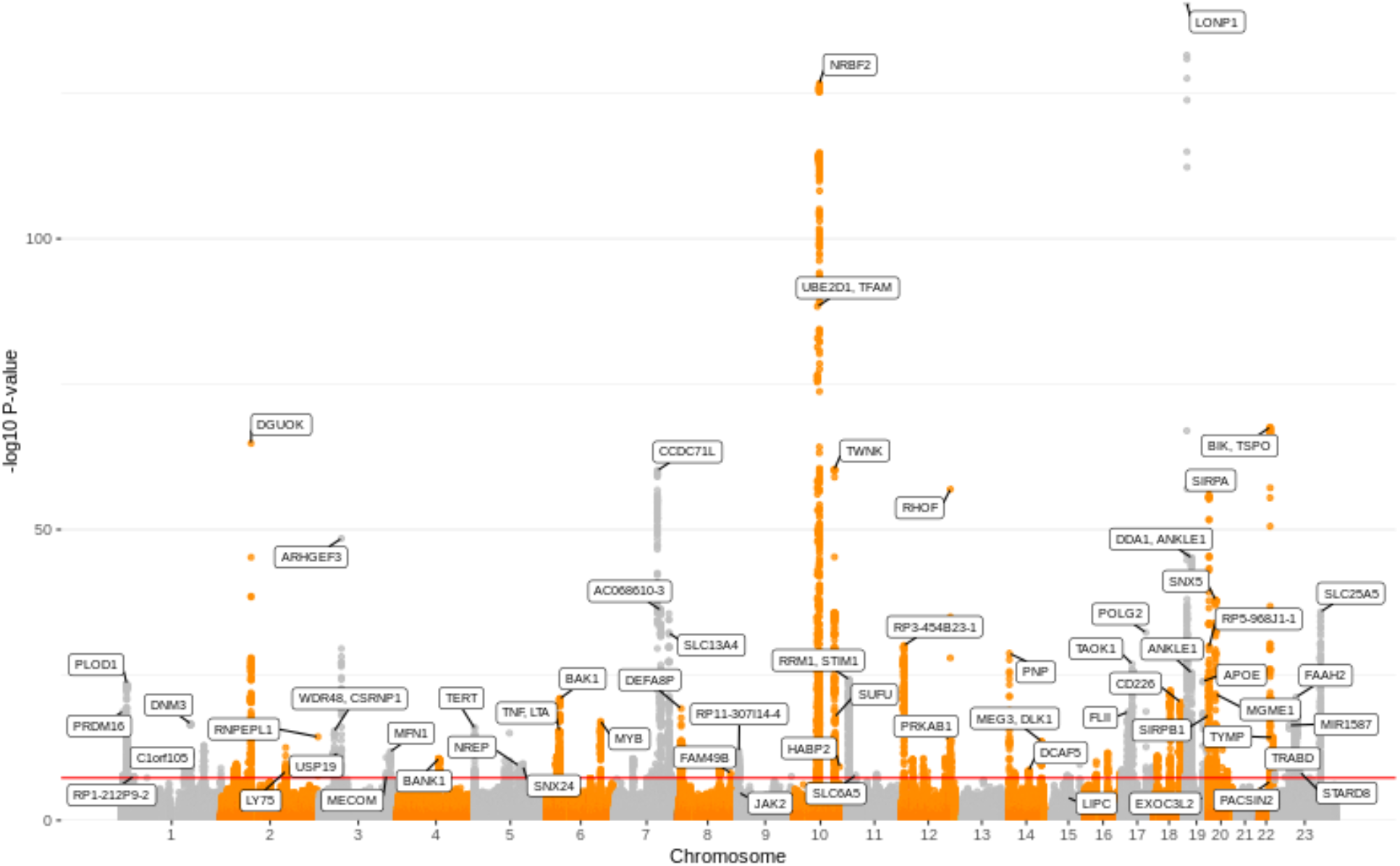
Manhattan plot of GWS loci from UKB-only analyses. Manhattan plot showing genome-wide significant loci for the UK Biobank-only analyses.

The most significant SNP associated with mtDNA-CN was a missense mutation in *LONP1* (p = 3.00×10^−141^), a gene that encodes a mitochondrial protease that can directly bind mtDNA, and has been shown to regulate *TFAM*, a transcription factor involved in mtDNA replication and repair (for review see Gibellini *et al.*)^64^.

Meta-analysis of the sex-stratified X chromosome results identified four loci significantly associated with mtDNA-CN, with directionality consistent across the male and female stratified analyses (Supplemental Table 7).

### Fine-mapping and Secondary Hits

To identify additional independent SNPs within novel loci whose effects were masked by the original significant SNP, as well as identify additional loci, we took two approaches. First, a conditional analysis adjusting for the top 93 SNPs from the initial (primary) GWAS run revealed 3 novel loci and 19 additional independent significant SNPs within existing loci. We also performed fine-mapping with susieR^42^ and discovered an additional 14 independent SNPs within existing loci. The majority of loci had only one 95% credible set of SNPs; further, twenty of the credible sets contained only one SNP. However, many of the credible sets contained greater than 50 SNPs after fine-mapping, and 12 of the 122 credible sets had a missense SNP as the SNP with the highest PIP in the set. Using these two methods, we identified in total 129 independent SNPs across 96 autosomal loci (Supplemental Figure 5), while susieR fine-mapping and conditional analyses for the X-chromosome loci did not reveal any additional secondary signals.

### Comparisons with Hägg et al. 2020

Out of the 50 loci reported in Hägg et al. 2020^33^, we replicate 38 loci in our cell-count adjusted analyses (Supplemental Table 8). As the two GWASs both use UK Biobank data, this replication is unsurprising. Out of the 12 loci that were not genome-wide significant in our cell-count adjusted analyses, 11 were significant when we did not adjust our mtDNA-CN metric for cell counts, suggesting that cell-type composition may be driving these signals. The current manuscript also reports 62 additional loci that are not in the Hägg et al. 2020 study. This is likely due to increased power, as the sample size used for the current analyses is nearly twice as large.

### Associations in Black Populations

Examining the 129 autosomal SNPs from the Whites-only analysis, 99 were available in the Blacks-only meta-analysis (n = 4770). After multiple testing correction, one of these SNPs was significant (rs73349121, *p* = 0.0001), 9 were nominally significant (*p* < 0.05, with 5 expected), and 58/99 had a direction of effect that was consistent with the Whites-only analyses (one-sided *p* = 0.04, Figure 2). Despite being under-powered, these results in the Blacks-only analyses provide evidence for similar genetic effects in a different ancestry group.

**Figure 2.**
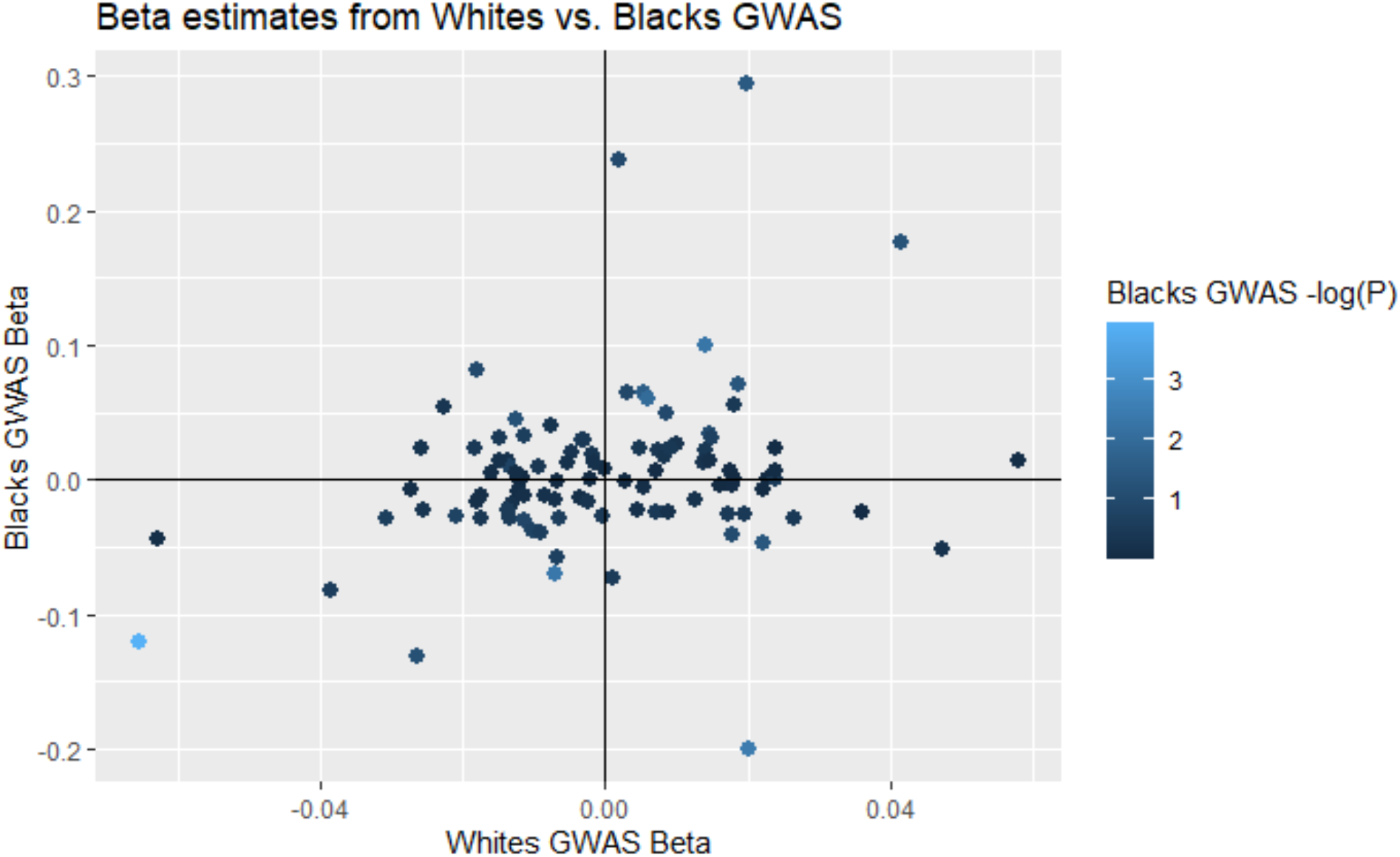
Scatterplot displaying effect size estimates between Whites and Blacks GWAS results for the 129 autosomal SNPs identified in the Whites analyses. Scatterplot showing comparison between effect size estimates for White and Black individuals. Color represents significance of effect for each locus in Blacks GWAS analyses.

### Gene Prioritization and Enrichment of mtDNA Depletion Syndrome Genes

We integrated results from three different gene prioritization and functional annotation methods (ANNOVAR^43^, COLOC^46^, and DEPICT^48^) so that loci with nonsynonymous variants in gene exons were prioritized first, with eQTL co-localization results considered second (Supplemental Table 9), and those from DEPICT (Supplemental Table 10) were considered last (Supplemental Figure 2). For 20 loci, multiple genes were assigned as analyses could not identify a single priority gene (Supplemental Table 11). As eQTLGen did not evaluate X chromosome variants, three of the four X-chromosome loci were assigned to the nearest gene. SLC25A5 was assigned to rs392020, as the second highest PIP SNP was an exonic nonsynonymous variant.

We noted the identification of a number of mtDNA depletion syndrome genes in the priority list and tested for enrichment of these known causal genes using a one-sided Fisher’s exact test. For this analysis, all genes for loci assigned to multiple genes were used, and genes for all primary and secondary loci were considered. Our gene prioritization approach identified 7 of 16 mtDNA depletion genes (Supplemental Table 12), consistent with a highly significant enrichment (one-sided *p* = 3.09 x 10^−15^).

### Gene Set Enrichment Analyses

To avoid bias from a single locus with multiple functionally related genes contributing to a false-positive signal, only one gene per unique locus was used, prioritizing genes assigned to primary loci. For loci with multiple assigned genes, one gene was randomly selected for testing. To test for robustness of gene set enrichment results, random selection was repeated 50 times, and only gene sets that were significantly enriched for at least 40 iterations were retained. In all, a total of 100 genes were utilized for GO term and KEGG pathway enrichment analyses. Using a Bonferroni-corrected p-value cutoff, 12 gene sets were significantly enriched for all 50 iterations, including mitochondrial nucleoid, mitochondrial DNA replication, and amyloid-beta clearance (Supplemental Table 13). No KEGG terms were significant across multiple iterations.

### MetaXcan Gene Expression Analysis

As a complementary approach to single-SNP analyses, we explored the associations between mtDNA-CN and predicted gene expression using MetaXcan^51^ MetaXcan incorporates multiple SNPs within a locus along with a reference eQTL dataset to generate predicted gene expression levels. As our study estimated mtDNA-CN derived from blood, we used whole blood gene expression eQTLs from the Gene-Tissue Expression (GTEx) consortium^65^ to predict gene expression in the UKB dataset. We identified 6,285 genes that had a predicted performance *p* < 0.05 (i.e., they had sufficient data to generate robust gene expression levels) and were tested for association with mtDNA-CN. Of these genes, 74 were significantly associated with mtDNA-CN (*p* < 7.95 x 10^−6^) (Figure 3, Supplemental Table 14), including 8 that were not identified through single-SNP analyses. Many of the significant genes have known mitochondrial functions, notably the mtDNA transcription factor *TFAM* (*p* = 1.09 x 10^−29^) and mitochondrial exonuclease *MGME1* (*p* = 5.87 x 10^−23^), genes known as causal for mtDNA depletion syndromes^54,55^. Additionally, *LONP1*, *MRPL43*, and *BAK1*, are all genes with known mitochondrial functions^66–68^. Bonferroni significant MetaXcan genes were used for gene enrichment analysis, finding enrichment for “nucleobase metabolic process” (*p* = 1.47 x 10^−4^) and “mitochondrial fusion” (*p* = 1.86 x 10^−4^) (Supplemental Figure 6).

**Figure 3.**
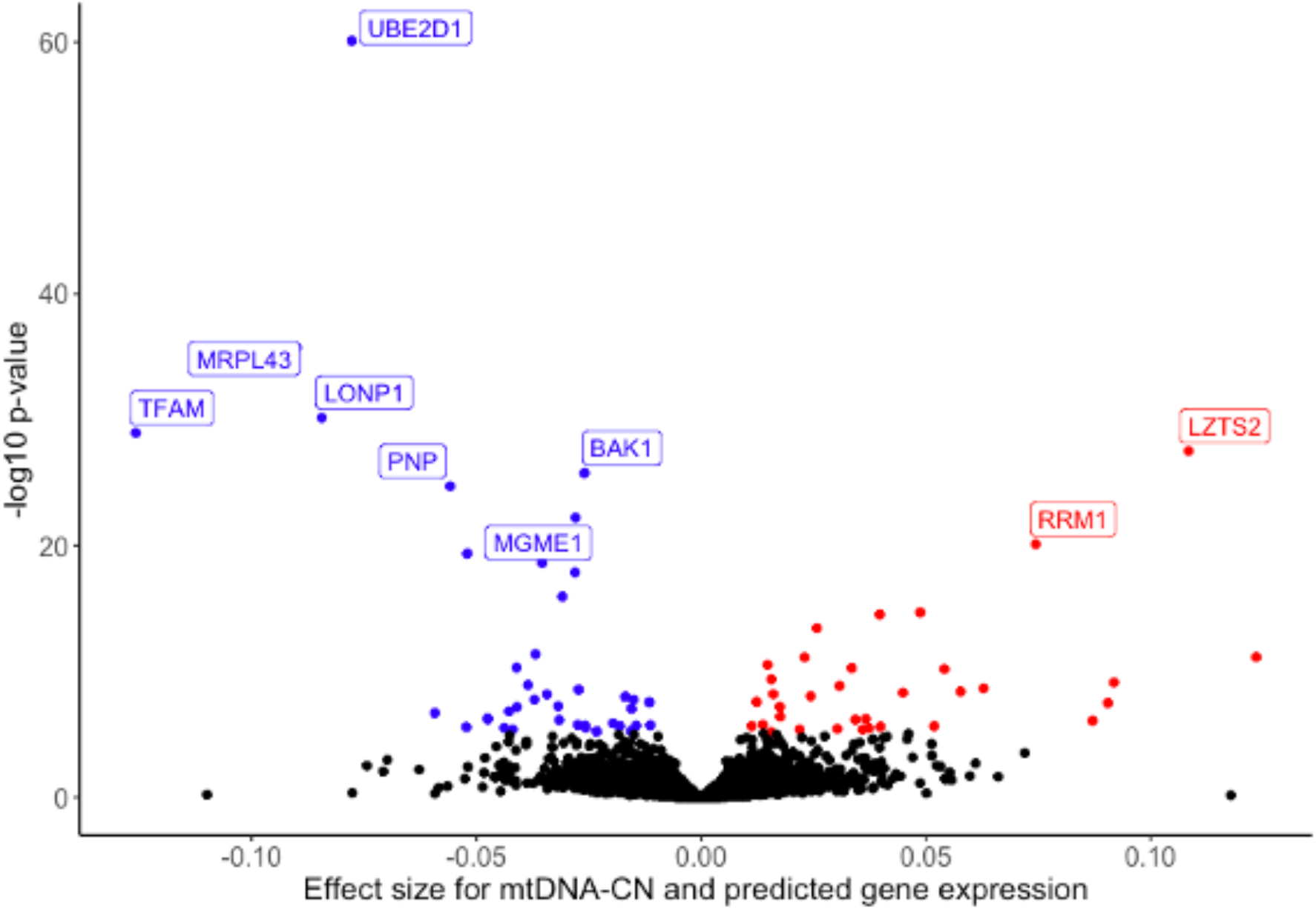
Volcano plot of genes whose predicted gene expression is significantly associated with mtDNA-CN. Volcano plot showing genes whose predicted gene expression is significantly associated with mtDNA-CN. Red indicates positive associations, blue indicates negative associations. Three genes (ARRDC1, EHMT1, PNPLA7) had extreme effect size estimates greater than 0.3 but were non-significant and removed from the plot for readability.

### PHEWAS-based SNP Clustering and Gene Set Enrichment

mtDNA-CN is associated with numerous quantitative and qualitative phenotypes, many of which are relevant to aging-related disease^3–5,9,10,13–16^. We hypothesized that this pleiotropy may reflect different underlying functional domains captured by mtDNA-CN, and may be reflected in GWAS-identified SNPs and their likely causal genes. To test this hypothesis, we used the UKB data to identify quantitative traits associated with mtDNA-CN and selected 41 highly significant, non-redundant traits to test for association with the mtDNA-CN GWAS SNPs (Supplemental Table 5, in PHEWAS = 1). We clustered SNPs using the trait effect size (beta) divided by the mtDNA-CN effect size estimate, so that all effects are standardized to the effect of the mtDNA-CN increasing allele for each locus. We identified 3 clusters of SNPs (Supplemental Table 15, Figure 4A), with cluster 1 containing SNPs in which the mtDNA-CN increasing allele is associated with decreased platelet count (PLT) (Figure 4B), increased mean platelet volume (MPV) (Figure 4C), and platelet distribution width (PDW) (Figure 4D), consistent with a role in platelet activation^69^. Cluster 2 is most strongly enriched for SNPs in which the mtDNA-CN increasing allele is associated with increased PLT, plateletcrit (PCT, a measure of total platelet mass), serum calcium (Figure 4E), serum phosphate, as well as decreased mean corpuscular volume (MCV) and mean spherical cellular volume (Figure 4F) (Supplemental Table 16). The cluster 2 phenotypes implicate megakaryocyte proliferation and proplatelet formation in addition to apoptosis and autophagy, and are supported by the genes identified for this cluster (megakaryocyte proliferation and proplatelet formation: *MYB*, *JAK2*^70^, apoptosis and autophagy: *BAK1*, *BCL2, TYMP*)^71^. Gene set enrichment analysis confirmed this, as cluster 2 genes are significantly enriched for extrinsic apoptosis signaling pathways in the absence of ligand (Supplemental Table 17). Cluster 3 did not yield any specific trait enrichment (all significant results reflected the strong enrichment observed in clusters 1-2); however, gene set enrichment for this cluster identified multiple mtDNA-related gene ontology terms, including mitochondrial DNA replication, gamma DNA polymerase complex, and mitochondrial nucleoid (Supplemental Table 18).

**Figure 4.**
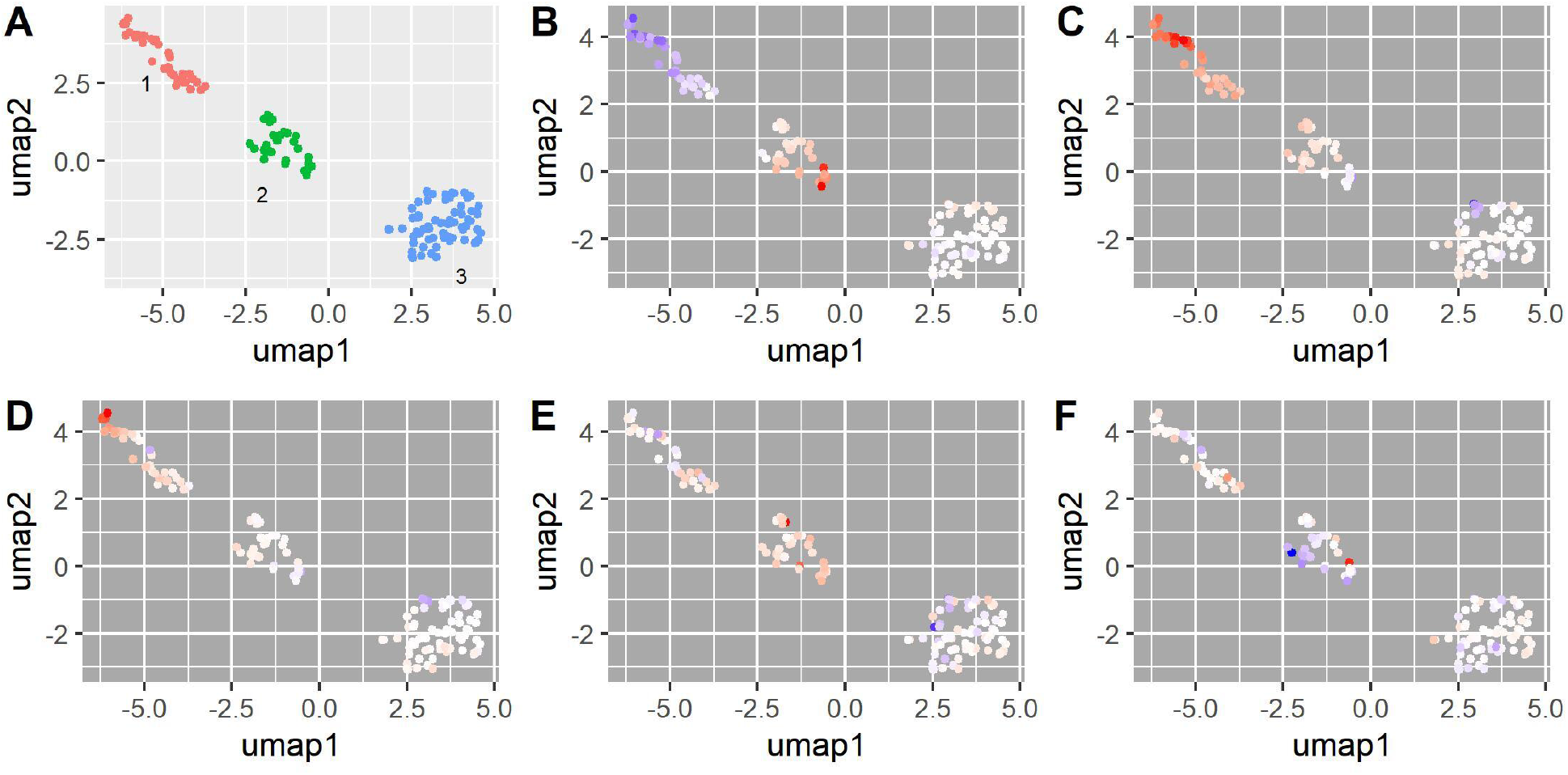
PHEWAS-based clustering of mtDNA-CN associated SNPs. UMAP clusters created from PHEWAS associations for mtDNA-CN associated SNPs. (A) Three clusters were identified as labeled in the panel; orange indicates no cluster. (B-F) SNPs are colored based on their effect estimate size, standardized to the effect on mtDNA-CN (red = positive, blue = negative estimates), for (B) platelet count, (C) mean platelet volume, (D) platelet distribution width, (E) serum calcium levels, (F) mean spherical cellular volume.

### Determination of causal associations between mitochondrial function and mtDNA-CN associated traits

The extensive pleiotropy and limited variance explained of nuclear DNA SNPs associated with mtDNA-CN (<1% of the variance in mtDNA-CN explained by GWS loci when predicted into the ARIC cohort) precludes the use of traditional Mendelian randomization (MR) approaches to establish causality between mtDNA-CN and the 41 identified mtDNA-CN associated traits. As alternative approach, we examined the association of mitochondrial SNPs with mtDNA-CN and the 41 traits, under the assumption that these SNPs can only act through alteration of mitochondrial function, and thus a significant association implies causality. Imputation and analyses of mitochondrial SNPs were run stratified by genotyping array (see Methods), and then meta-analyzed using inverse-variance weighting. After multi-test correction (*p* < 9.5×10^−6^), we identified 45 SNPs associated with 1 or more of the traits, ranging from 1 to 6 traits per SNP (Supplementary Table 19). To identify independent effects, we first identified the most significantly associated trait for each SNP, highlighting 4 traits (aspartate aminotransferase, creatinine, MCV, PCT) in which to run susieR to identify independent credible sets. We identified 6 independent effects across the four traits, with MCV credible set 4 and platelet credible set 1 representing the same effect. We note that 2 of the SNPs are in moderately high LD (MT73A_G and MT7028C_T, r^2^=0.67), however, conditional analyses demonstrate that MT73A_G is associated with creatinine, and not MCV, and the reverse is seen for MT7028C_T (Supplemental Table 20). Leveraging the haploid nature of the mitochondrial genome, we selected the directly genotyped SNP with the highest PIP from each credible set (Supplemental Table 21), and identified 8 haplotypes with MAF > 0.005 (Supplemental Table 22). Comparing linear regression models with and without the haplotypes in the model, we identify 14 traits nominally associated, and 9 traits significantly associated after Bonferroni correction, with mtDNA genetic variation (Supplemental Table 23, Figure 5). These results causally implicate mitochondrial function in a variety of cell related traits (MCV, MSCV, MPV, PCT, Platelet), kidney function (creatinine), liver function (aspartate and alanine aminotransferases) and mtDNA-CN.

**Figure 5.**
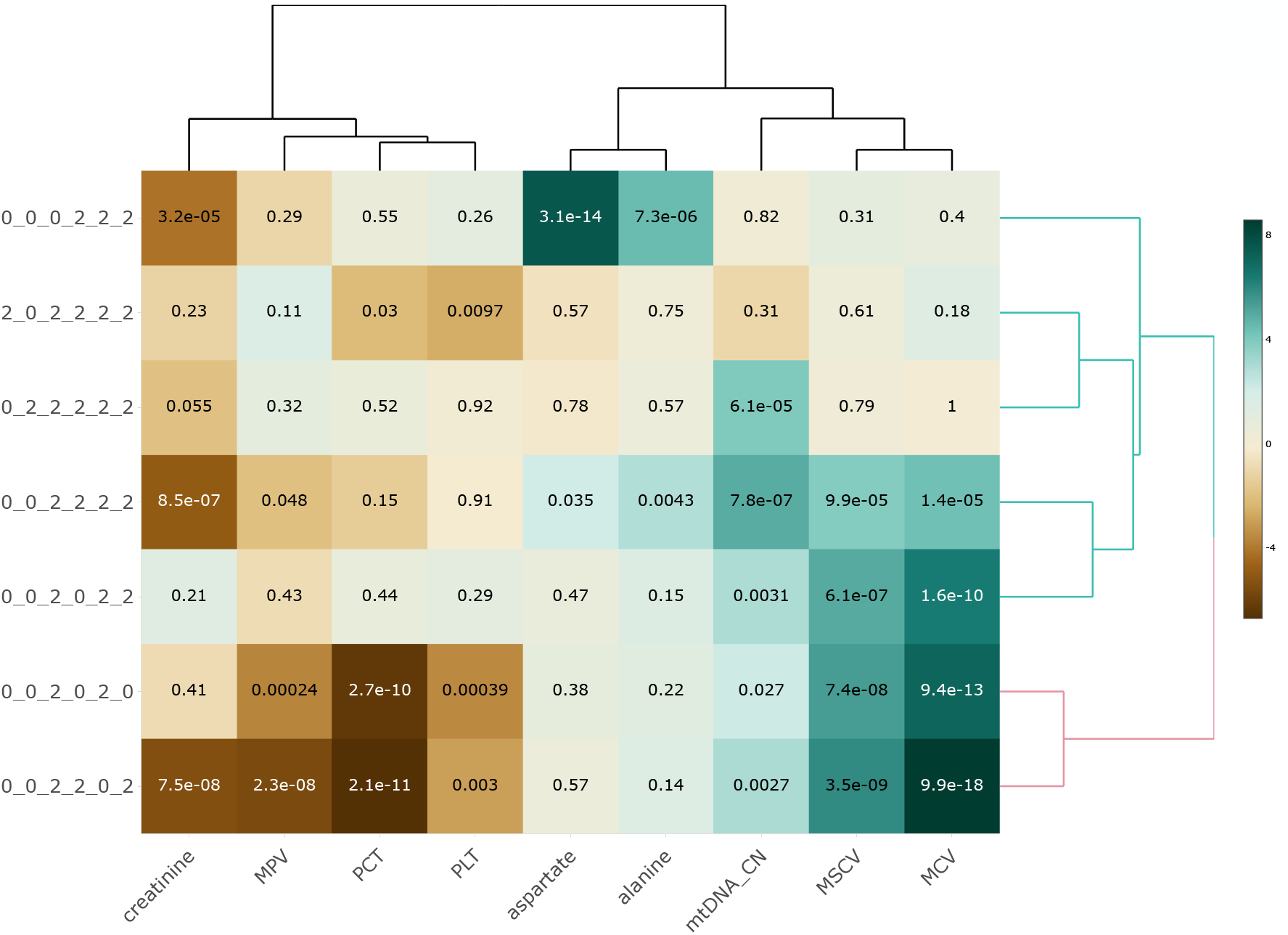
Associations between mtDNA-CN associated phenotypes and mitochondrial haplotypes. Mitochondrial haplotypes are significantly associated with mtDNA-CN associated traits, implying causal relationships between mitochondrial function and traits of interest. Haplotypes are notated in the following format: MT73_MT12612_MT7028_MT10238_MT13617_MT15257.

### Association of mitochondrial function with mortality

We have previously shown that mtDNA-CN is associated with overall mortality^9^. As above, we collectively tested the mitochondrial haplotypes for association with mortality not due to external causes (e.g., no accidents, falls, see Methods; n = 24,622, median follow-up time = 4318 days), and found a nominally significant association with overall mortality (*p* = 0.044, Figure 6). Given the conflicting reports between increased mitochondrial function and both increased and decreased cancer risk^16,72,73^, we looked separately at cancer (n = 13,231) and non-cancer mortality (n = 11,391). While there was no association with cancer mortality (*p* = 0.74), we saw a highly significant association with non-cancer mortality (*p* = 6.56×10^−4^).

**Figure 6.**
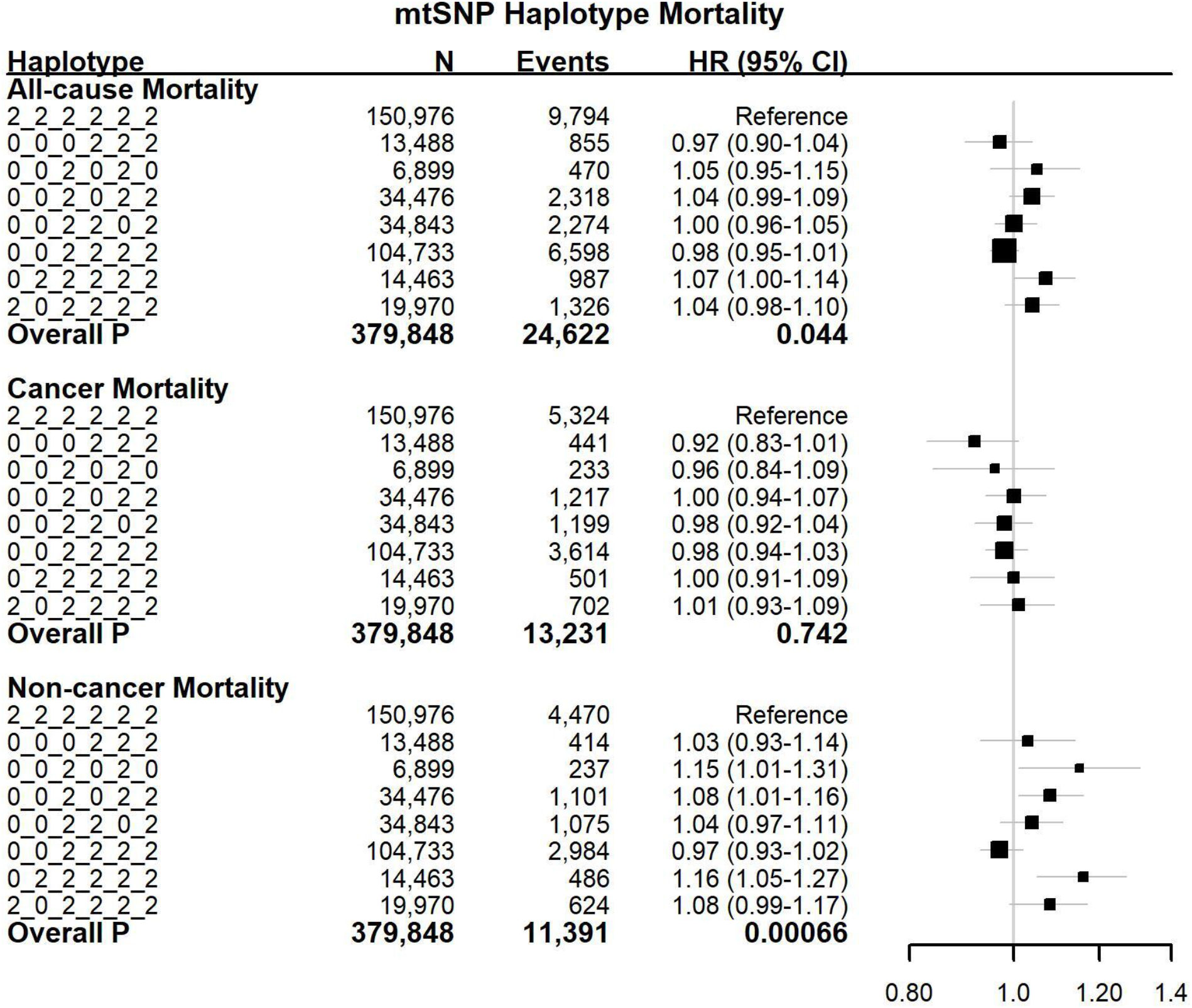
Associations between mitochondrial DNA haplotypes and morbidity (overall, cancer, and noncancer) Mitochondrial haplotypes are significantly associated with overall morbidity, in particular, non-cancer mortality. Haplotypes are notated in the following format: MT73_ MT7028_MT10238_ MT12612_MT13617_MT15257.

## Discussion

We conducted a GWAS for mtDNA-CN using 465,809 individuals from the CHARGE consortium and the UKB. We report 133 independent signals originating from 100 loci, the majority of which were not identified in previous studies. Examining our GWS SNPs in a Black population, we observed a concordant signal, suggesting that the genetic etiology of mtDNA-CN may be broadly similar across populations. Using several functional follow-up methods, genes were assigned for each identified independent hit and significant enrichment was observed for genes involved in mitochondrial DNA metabolism, homeostasis, cell activation, and amyloid-beta clearance. In total, we assigned 128 unique genes to independent GWAS signals associated with mtDNA-CN. We also identified 8 additional genes whose predicted gene expression is associated with mtDNA-CN that could not be mapped back to GWS loci. Finally, using a clustering approach based on SNP associations with various mtDNA-CN associated phenotypes, we were able to functionally categorize SNPs, providing insight into biological pathways that impact mtDNA-CN.

We note that during the preparation of this manuscript, a GWAS for mtDNA-CN performed in 295,150 unrelated individuals from the UK Biobank was published, which reported 50 genome-wide significant regions^33^. Within our GWAS, we replicate 38 of these 50 genome-wide significant loci in our cell count corrected analyses. An additional 11 out of the remaining 12 loci are genome-wide significant when we do not adjust mtDNA-CN for cell count. While Hagg et. al adjust for cell type composition, this difference suggests that their adjustment may not be fully capturing the effects of cell counts. Additionally, our analyses report 59 additional loci that are not observed in the previous paper, largely due to the increased power of our study.

We were able to identify a substantial proportion of the genes involved in mtDNA depletion syndromes (7/16, *p* = 3.09 x 10^−15^ for enrichment), including *TWNK*, *TFAM*, *DGUOK*, *MGME1*, *RRM2B, TYMP,* and *POLG*. mtDNA depletion syndromes can be broken down into 5 subtypes based on their constellation of phenotypes^74^, and with the exception of cardiomyopathic subtypes (associated with mutations in *AGK* and *SLC25A4*), we were able to identify at least 1 gene from the other 4 subtypes, suggesting that our mtDNA-CN measurement in blood-derived DNA can identify genes widely relevant to non-blood phenotypes. This finding is consistent with a large body of work showing that mtDNA-CN measured in blood is associated with numerous aging-related phenotypes for which the primary tissue of interest is not blood (e.g. chronic kidney disease^13^, heart failure^11^, and diabetes^75^). Also consistent with this finding is recent work demonstrating that mtDNA-CN measured in blood is associated with mtRNA expression across numerous non-blood tissues, suggesting a link between mitochondrial function measured in blood and other tissues^76^.

In addition to identifying the mtDNA depletion syndrome genes directly linked to mitochondrial DNA metabolic processes, DNA replication, and genome maintenance, we also identify genes which play a role in mitochondrial function. The top GWAS hit is a missense mutation in *LONP1*, which encodes a mitochondrial protease that has been shown to cause mitochondrial cytopathy and reduced respiratory chain activity^77,78^. Interestingly, this missense mutation was recently found to be associated with mitochondrial tRNA methylation levels^79^. Additional genes known to impact mitochondrial function include *MFN1*, which encodes a mediator of mitochondrial fusion^80,81^, *STMP1*, which plays a role in mitochondrial respiration^82^, and *MRPS35*, which encodes a ribosomal protein involved in protein synthesis in the mitochondrion^83,84^.

Using a combination of gene-based tests and gene prioritization using functional annotation, pathway analyses reveal enrichment for numerous mitochondrial related pathways, as well as those involved in regulation of cell differentiation (*p* < 1.08 x 10^−5^), homeostatic processes (*p* < 3.77 x 10^−6^), and cellular response to stress (*p* < 3.49 x 10^−6^) (Supplemental Table 13). These results provide additional evidence for the broad role played by mitochondria in numerous aspects of cellular function. Of particular interest, the GO term for amyloid beta is significantly enriched, reinforcing a link between mtDNA-CN and neurodegenerative disease^85–87^. Previous work from our lab using the UKB has shown that increased mtDNA-CN is associated with lower rates of prevalent neurodegenerative disease, and is predictive for decreased risk of incident neurodegenerative disease^76^. mtDNA-CN is also known to be decreased in the frontal cortex of Alzheimer’s disease (AD) patients^88^. Interestingly, the four GWAS-identified genes driving the enrichment for amyloid-beta clearance are all related to regulation of lipid levels, and lipid homeostasis within the brain is known to play an important role in Alzheimer’s disease^89^. *APOE*, one of the most well-known risk genes for Alzheimer’s disease, is a cholesterol carrier involved in lipid transport, and the ApoE-ɛ4 isoform involved in AD pathogenesis is associated with mitochondrial dysfunction and oxidative distress in the human brain^90^; *CD36* is a platelet glycoprotein which mediates the response to amyloid-beta accumulation^91^; *LDLR* is a low-density lipoprotein receptor associated with AD^92^; and *ABCA7* is a phospholipid transporter^93^. *ABCA7* loss of function variants are enriched in both AD and Parkinson’s disease (PD) patients^94^, suggesting a broad role across neurodegenerative diseases.

Given the integral role of mitochondria in cellular function, from ATP formation and energy production, signaling through reactive oxygen species, and apoptosis mediation, there is a strong basis to *a priori* assume that genetic variants associated with mtDNA-CN are likely to be highly pleiotropic. Indeed, mtDNA-CN itself is associated with numerous phenotypes (Supplemental Table 5). Through our PHEWAS-based clustering approach using 41 mtDNA-CN associated phenotypes, we uncovered phenotypic associations between three distinct clusters of GWS mtDNA-CN associated SNPs. Cluster 1 was characterized by increased MPV, PDW, and decreased PLT (note that measured MPV and PLT are generally inversely correlated to maintain hemostasis), which are the hallmarks of platelet activation^69^. The link between platelets and mtDNA-CN has typically revolved around platelet count, as platelets have functional mitochondria, but do not have a nucleus. Given that the mtDNA-CN measurement is the ratio between mtDNA and nuclear DNA, increased platelets, all else being equal, would directly equate with increased mtDNA-CN. We note that the mtDNA-CN metric used in this GWAS was adjusted for platelet count, likely increasing the ability to detect variants that impact mtDNA-CN through increased platelet activation. Examining the genes within this cluster suggests roles for actin formation and regulation (*TPM4*, *PACSIN2*)^95,96^ and vesicular transport and endocytic trafficking (*DNM3*, *EHD3*)^97,98^ in platelet activation.

Cluster 2 is most strongly enriched for SNPs in which the mtDNA-CN increasing allele is associated with increased PLT/PCT and serum calcium/phosphate. Examining the genes assigned to the cluster, we implicate megakaryocyte proliferation and proplatelet formation (*MYB*, *JAK2*)^70^, and apoptosis and autophagy (*BAK1*, *BCL2, TYMP*)^71^. Megakaryocytes are used to form proplatelets, and the process includes an important role for both intra- and extracellular calcium levels^99^. A role for apoptosis, and specifically *BCL2*, in proplatelet formation and platelet release has been suggested^100,101^, however work in mice has suggested that apoptosis does not play a direct role in these processes^102^. Nevertheless, apoptosis is important for platelet lifespan^103^.

Cluster 3 was particularly challenging to interpret, given that no particular phenotype was enriched relative to the non-cluster 3 SNPs. We note that this cluster appeared to be enriched for the mtDNA depletion syndrome genes, containing 6/7 genes identified in the GWAS, and significantly enriched for GO Terms related mitochondrial DNA. Additionally, genes in cluster 3 were significantly enriched for low-density lipoprotein particle binding, suggesting a role for lipid homeostasis. Closer inspection of cluster 3 genes reveals a number of genes known to be associated with lipid levels (*LIPC*, *CETP*, *LDLR*, *APOE*). While lipids play a role in both energy metabolism (largely through fatty acids) and cellular membrane formation, a link to mtDNA-CN and/or mitochondrial function is not well-established.

A strong rationale for the study of mtDNA-CN is the underlying assumption that it reflects mitochondrial function and is readily measured, often from existing data. A serious complication to the interpretation of the role of mitochondrial function in various traits has been use of blood-derived measurements, which can be confounded by differences in cell counts across individuals. Mendelian randomization has been widely used to infer causality between traits (e.g. LDL and CAD)^104^, but is only robust under conditions of little to no pleiotropy^105^ and its power is a function of variance explained. For mtDNA-CN, the extensive pleiotropy and small amount of variance explained of GWS variants (<1%) prevents the use of traditional MR approaches. As an alternative approach, we analyzed associations between mitochondrial DNA variants and mtDNA-CN associated phenotypes. Presumably, variants located on the mitochondrial genome are only able to modify phenotypes through modulating mitochondrial function, allowing for causal inference. Our analyses revealed significant relationships between mitochondrial variants and creatinine, aspartate aminotransferase, MCV, and PCT. Creatinine and aspartate aminotransferase are markers of kidney and liver function respectively, and supporting these findings, mtDNA-CN has been linked to both chronic kidney disease^13^ and non-alcoholic fatty liver disease^106^. We also find a highly significant association between mitochondrial variation and non-cancer mortality, adding evidence for a causal relationship to previous findings showing mtDNA-CN is associated with all-cause mortality^9^.

Several limitations should be noted. First, despite the large sample size and numerous loci identified, we are likely missing a great deal of the true signal, as our SNP heritability estimates through SumHer and BOLT-LMM were 7.0% and 7.4% respectively, while previous studies have estimated mtDNA-CN heritability to be 65%^23^. Finally, while we have adjusted our mtDNA-CN metric for a variety of confounders, it is important to note that mtDNA-CN can be influenced by a variety of environmental factors including smoking^107^ and drugs, which have not been adjusted for in these analyses.

In summary, we performed the largest-to-date GWAS for mtDNA-CN, including almost 500,000 individuals. We identified three distinct groups of SNPs associated with mtDNA-CN that are related to platelet activation, megakaryocyte formation and apoptotic processes, and showed clear enrichment for genes involved in mtDNA depletion and nucleotide regulation. Additionally, we find that mitochondrial variants are significantly associated with creatinine, aspartate aminotransferase, MCV, and PCT, implying a causal relationship between mitochondrial function and these phenotypes. Finally, we provide strong evidence that mitochondrial function is causal for non-cancer mortality. Given the role of mtDNA-CN, and, by proxy, mitochondrial function in aging-related disease, this work begins to unravel the many varied underlying mechanisms through which mitochondrial function impacts human health.

## Supplemental Data

Supplemental Data include six supplemental figures, twenty-two supplemental tables, and supplemental methods.

## Acknowledgements

This work was supported by National Heart, Lung and Blood Institute, National Institutes of Health (NIH) grants R01HL13573 and R01HL144569 (RJL, SYY, CAC, DEA), NIH grant P01-AG027734 (GA, YK, NB, AB) and the National Center for Advancing Translational Sciences, NIH, through Grant KL2TR002490 (LMR). The content is solely the responsibility of the authors and does not necessarily represent the official views of the NIH. LMR was also funded by T32 HL129982.

This research was conducted using data from the Genotype-Tissue Expression (GTEx) project (dbGaP accession: phs000424.**v8**.p2). The GTEx project was supported by the Common Fund of the Office of the Director of the National Institutes of Health, and by NCI, NHGRI, NHLBI, NIDA, NIMH, and NINDS.

This research was also conducted using the UK Biobank Resource under Application Number 17731.

The Atherosclerosis Risk in Communities study (dbGaP accession: phs000280.**v7**.p1) has been funded in whole or in part with Federal funds from the National Heart, Lung, and Blood Institute, National Institutes of Health, Department of Health and Human Services (contract numbers HSN268201700001I, HHSN268201700002I, HHSN268201700003I, HHSN268201700004I and HHSN268201700005I), R01HL087641, R01HL059367 and R01HL086694; National Human Genome Research Institute contract U01HG004402; and National Institutes of Health contract HHSN268200625226C. Funding support for “Building on GWAS for NHLBI diseases: the U.S. CHARGE consortium” was provided by the NIH through the American Recovery and Reinvestment Act of 2009 (ARRA) (5RC2HL102419). Sequencing was carried out at the Baylor College of Medicine Human Genome Sequencing Center and supported by the National Human Genome Research Institute grants U54 HG003273 and UM1 HG008898. The authors thank the staff and participants of the ARIC study for their important contributions. Infrastructure was partly supported by Grant Number UL1RR025005, a component of the National Institutes of Health and NIH Roadmap for Medical Research.

This CHS research was supported by NHLBI contracts HHSN268201200036C, HHSN268200800007C, HHSN268201800001C, N01HC55222, N01HC85079, N01HC85080, N01HC85081, N01HC85082, N01HC85083, N01HC85086, 75N92021D00006; and NHLBI grants U01HL080295, R01HL087652, R01HL105756, R01HL103612, R01HL120393, and U01HL130114 with additional contribution from the National Institute of Neurological Disorders and Stroke (NINDS). Additional support was provided through R01AG023629 from the National Institute on Aging (NIA). A full list of principal CHS investigators and institutions can be found at CHS-NHLBI.org. The provision of genotyping data was supported in part by the National Center for Advancing Translational Sciences, CTSI grant UL1TR001881, and the National Institute of Diabetes and Digestive and Kidney Disease Diabetes Research Center (DRC) grant DK063491 to the Southern California Diabetes Endocrinology Research Center.

The FHS phenotype-genotype analyses were supported by the National Institute of Aging (U34AG051418). This research was conducted in part using data and resources from the Framingham Heart Study of the National Heart Lung and Blood Institute of the National Institutes of Health and Boston University School of Medicine. This work was partially supported by the National Heart, Lung and Blood Institute’s Framingham Heart Study (Contract No. N01-HC-25195, HHSN268201500001) and its contract with Affymetrix, Inc for genotyping services (Contract No. N02-HL-6-4278). Genotyping, quality control and calling of the Illumina HumanExome BeadChip in the Framingham Heart Study was supported by funding from the National Heart, Lung and Blood Institute Division of Intramural Research (Daniel Levy and Christopher J. O’Donnell, Principle Investigators). The authors thank the participants for their dedication to the study. The authors are pleased to acknowledge that the computational work reported on in this paper was performed on the Shared Computing Cluster which is administered by Boston University’s Research Computing Services. URL: www.bu.edu/tech/support/research/.

MESA and the MESA SHARe projects (dbGaP accession: phs000209.**v13**.p3) are conducted and supported by the National Heart, Lung, and Blood Institute (NHLBI) in collaboration with MESA investigators. Support for MESA is provided by contracts 75N92020D00001, HHSN268201500003I, N01-HC-95159, 75N92020D00005, N01-HC-95160, 75N92020D00002, N01-HC-95161, 75N92020D00003, N01-HC-95162, 75N92020D00006, N01-HC-95163, 75N92020D00004, N01-HC-95164, 75N92020D00007, N01-HC-95165, N01-HC-95166, N01-HC-95167, N01-HC-95168, N01-HC-95169, UL1-TR-000040, UL1-TR-001079, and UL1-TR-001420. Funding for SHARe genotyping was provided by NHLBI Contract N02-HL-64278. Genotyping was performed at Affymetrix (Santa Clara, California, USA) and the Broad Institute of Harvard and MIT (Boston, Massachusetts, USA) using the Affymetrix Genome-Wide Human SNP Array 6.0. The provision of genotyping data was supported in part by the National Center for Advancing Translational Sciences, CTSI grant UL1TR001881, and the National Institute of Diabetes and Digestive and Kidney Disease Diabetes Research Center (DRC) grant DK063491 to the Southern California Diabetes Endocrinology Research Center.

ROS/MAP is supported by the Translational Genomics Research Institute and National Institute on Aging (NIA) through grants U01AG46152, U01AG61256, P30AG10161, R01AG17917, RF1AG15819, R01AG30146.

SHIP and SHIP-TREND are part of the Community Medicine Research net of the University of Greifswald, Germany, which is funded by the Federal Ministry of Education and Research (grants no. 01ZZ9603, 01ZZ0103, and 01ZZ0403), the Ministry of Cultural Affairs as well as the Social Ministry of the Federal State of Mecklenburg-West Pomerania, and the network ‘Greifswald Approach to Individualized Medicine (GANI_MED)’ funded by the Federal Ministry of Education and Research (grant 03IS2061A).

This research was funded in whole, or in part, by the Wellcome Trust [Grant ref: 217065/Z/19/Z]. For the purpose of Open Access, the author has applied a CC BY public copyright license to any Author Accepted Manuscript version arising from this submission.

## Web Resources

REVIGO was accessed at http://revigo.irb.hr/.

## Data and Code Availability

All data used in this manuscript is available through either the UKBiobank and CHARGE consortiums. Code and scripts are available in a zipped file at https://www.arkinglab.org/resources/.

## References

1. Wallace, D.C. (1992). Diseases of the Mitochondrial Dna. Annual Review of Biochemistry 61, 1175–1212.

2. Vakifahmetoglu-Norberg, H., Ouchida, A.T., and Norberg, E. (2017). The role of mitochondria in metabolism and cell death. Biochem. Biophys. Res. Commun. 482, 426–431.

3. Dai, D.-F., Rabinovitch, P.S., and Ungvari, Z. (2012). Mitochondria and cardiovascular aging. Circ. Res. 110, 1109–1124.

4. Cui, H., Kong, Y., and Zhang, H. (2012). Oxidative stress, mitochondrial dysfunction, and aging. J Signal Transduct 2012, 646354.

5. Herst, P.M., Rowe, M.R., Carson, G.M., and Berridge, M.V. (2017). Functional Mitochondria in Health and Disease. Front Endocrinol (Lausanne) 8, 296.

6. Liu, C.-S., Tsai, C.-S., Kuo, C.-L., Chen, H.-W., Lii, C.-K., Ma, Y.-S., and Wei, Y.-H. (2003). Oxidative stress-related alteration of the copy number of mitochondrial DNA in human leukocytes. Free Radic Res 37, 1307–1317.

7. Guha, M., and Avadhani, N.G. (2013). Mitochondrial retrograde signaling at the crossroads of tumor bioenergetics, genetics and epigenetics. Mitochondrion 13, 577–591.

8. Malik, A.N., and Czajka, A. (2013). Is mitochondrial DNA content a potential biomarker of mitochondrial dysfunction? Mitochondrion 13, 481–492.

9. Ashar, F.N., Moes, A., Moore, A.Z., Grove, M.L., Chaves, P.H.M., Coresh, J., Newman, A.B., Matteini, A.M., Bandeen-Roche, K., Boerwinkle, E., et al. (2015). Association of mitochondrial DNA levels with frailty and all-cause mortality. J. Mol. Med. 93, 177–186.

10. Ashar, F.N., Zhang, Y., Longchamps, R.J., Lane, J., Moes, A., Grove, M.L., Mychaleckyj, J.C., Taylor, K.D., Coresh, J., Rotter, J.I., et al. (2017). Association of Mitochondrial DNA Copy Number With Cardiovascular Disease. JAMA Cardiol 2, 1247–1255.

11. Hong, Y.S., Longchamps, R.J., Zhao, D., Castellani, C.A., Loehr, L.R., Chang, P.P., Matsushita, K., Grove, M.L., Boerwinkle, E., Arking, D.E., et al. (2020). Mitochondrial DNA Copy Number and Incident Heart Failure: The Atherosclerosis Risk in Communities (ARIC) Study. Circulation 141, 1823–1825.

12. Zhao, D., Bartz, T.M., Sotoodehnia, N., Post, W.S., Heckbert, S.R., Alonso, A., Longchamps, R.J., Castellani, C.A., Hong, Y.S., Rotter, J.I., et al. (2020). Mitochondrial DNA copy number and incident atrial fibrillation. BMC Medicine 18, 246.

13. Tin, A., Grams, M.E., Ashar, F.N., Lane, J.A., Rosenberg, A.Z., Grove, M.L., Boerwinkle, E., Selvin, E., Coresh, J., Pankratz, N., et al. (2016). Association between Mitochondrial DNA Copy Number in Peripheral Blood and Incident CKD in the Atherosclerosis Risk in Communities Study. J Am Soc Nephrol 27, 2467–2473.

14. Pyle, A., Anugrha, H., Kurzawa-Akanbi, M., Yarnall, A., Burn, D., and Hudson, G. (2016). Reduced mitochondrial DNA copy number is a biomarker of Parkinson’s disease. Neurobiol. Aging 38, 216.e7-216.e10.

15. Wei, W., Keogh, M.J., Wilson, I., Coxhead, J., Ryan, S., Rollinson, S., Griffin, H., Kurzawa-Akanbi, M., Santibanez-Koref, M., Talbot, K., et al. (2017). Mitochondrial DNA point mutations and relative copy number in 1363 disease and control human brains. Acta Neuropathologica Communications 5, 13.

16. Reznik, E., Miller, M.L., Şenbabaoğlu, Y., Riaz, N., Sarungbam, J., Tickoo, S.K., Al-Ahmadie, H.A., Lee, W., Seshan, V.E., Hakimi, A.A., et al. (2016). Mitochondrial DNA copy number variation across human cancers. ELife 5, e10769.

17. Hurtado-Roca, Y., Ledesma, M., Gonzalez-Lazaro, M., Moreno-Loshuertos, R., Fernandez-Silva, P., Enriquez, J.A., and Laclaustra, M. (2016). Adjusting MtDNA Quantification in Whole Blood for Peripheral Blood Platelet and Leukocyte Counts. PLOS ONE 11, e0163770.

18. Knez, J., Winckelmans, E., Plusquin, M., Thijs, L., Cauwenberghs, N., Gu, Y., Staessen, J.A., Nawrot, T.S., and Kuznetsova, T. (2016). Correlates of Peripheral Blood Mitochondrial DNA Content in a General Population. Am J Epidemiol 183, 138–146.

19. Kumar, P., Efstathopoulos, P., Millischer, V., Olsson, E., Wei, Y.B., Brüstle, O., Schalling, M., Villaescusa, J.C., Ösby, U., and Lavebratt, C. (2018). Mitochondrial DNA copy number is associated with psychosis severity and anti-psychotic treatment. Sci Rep 8, 12743.

20. Urata, M., Koga-Wada, Y., Kayamori, Y., and Kang, D. (2008). Platelet contamination causes large variation as well as overestimation of mitochondrial DNA content of peripheral blood mononuclear cells. Ann Clin Biochem 45, 513–514.

21. Clay Montier, L.L., Deng, J.J., and Bai, Y. (2009). Number matters: control of mammalian mitochondrial DNA copy number. J Genet Genomics 36, 125–131.

22. Tang, Y., Schon, E.A., Wilichowski, E., Vazquez-Memije, M.E., Davidson, E., and King, M.P. (2000). Rearrangements of Human Mitochondrial DNA (mtDNA): New Insights into the Regulation of mtDNA Copy Number and Gene Expression. Mol Biol Cell 11, 1471–1485.

23. Xing, J., Chen, M., Wood, C.G., Lin, J., Spitz, M.R., Ma, J., Amos, C.I., Shields, P.G., Benowitz, N.L., Gu, J., et al. (2008). Mitochondrial DNA content: its genetic heritability and association with renal cell carcinoma. J. Natl. Cancer Inst. 100, 1104–1112.

24. Carling, P.J., Cree, L.M., and Chinnery, P.F. (2011). The implications of mitochondrial DNA copy number regulation during embryogenesis. Mitochondrion 11, 686–692.

25. Harvey, A., Gibson, T., Lonergan, T., and Brenner, C. (2011). Dynamic regulation of mitochondrial function in preimplantation embryos and embryonic stem cells. Mitochondrion 11, 829–838.

26. Copeland, W.C. (2014). Defects of Mitochondrial DNA Replication. J Child Neurol 29, 1216–1224.

27. Mandel, H., Szargel, R., Labay, V., Elpeleg, O., Saada, A., Shalata, A., Anbinder, Y., Berkowitz, D., Hartman, C., Barak, M., et al. (2001). The deoxyguanosine kinase gene is mutated in individuals with depleted hepatocerebral mitochondrial DNA. Nat. Genet. 29, 337–341.

28. Wang, L., Limongelli, A., Vila, M.R., Carrara, F., Zeviani, M., and Eriksson, S. (2005). Molecular insight into mitochondrial DNA depletion syndrome in two patients with novel mutations in the deoxyguanosine kinase and thymidine kinase 2 genes. Mol. Genet. Metab. 84, 75–82.

29. Rusecka, J., Kaliszewska, M., Bartnik, E., and Tońska, K. (2018). Nuclear genes involved in mitochondrial diseases caused by instability of mitochondrial DNA. J Appl Genet 59, 43–57.

30. Cai, N., Li, Y., Chang, S., Liang, J., Lin, C., Zhang, X., Liang, L., Hu, J., Chan, W., Kendler, K.S., et al. (2015). Genetic Control over mtDNA and Its Relationship to Major Depressive Disorder. Curr Biol 25, 3170–3177.

31. Workalemahu, T., Enquobahrie, D.A., Tadesse, M.G., Hevner, K., Gelaye, B., Sanchez, S., and Williams, M.A. (2017). Genetic Variations Related to Maternal Whole Blood Mitochondrial DNA Copy Number: A Genome-Wide and Candidate Gene Study. J Matern Fetal Neonatal Med 30, 2433–2439.

32. Guyatt, A.L., Brennan, R.R., Burrows, K., Guthrie, P.A.I., Ascione, R., Ring, S.M., Gaunt, T.R., Pyle, A., Cordell, H.J., Lawlor, D.A., et al. (2019). A genome-wide association study of mitochondrial DNA copy number in two population-based cohorts. Human Genomics 13, 6.

33. Hägg, S., Jylhävä, J., Wang, Y., Czene, K., and Grassmann, F. (2020). Deciphering the genetic and epidemiological landscape of mitochondrial DNA abundance. Hum Genet.

34. Psaty Bruce M., O’Donnell Christopher J., Gudnason Vilmundur, Lunetta Kathryn L., Folsom Aaron R., Rotter Jerome I., Uitterlinden André G., Harris Tamara B., Witteman Jacqueline C.M., and Boerwinkle Eric (2009). Cohorts for Heart and Aging Research in Genomic Epidemiology (CHARGE) Consortium. Circulation: Cardiovascular Genetics 2, 73–80.

35. Bycroft, C., Freeman, C., Petkova, D., Band, G., Elliott, L.T., Sharp, K., Motyer, A., Vukcevic, D., Delaneau, O., O’Connell, J., et al. (2018). The UK Biobank resource with deep phenotyping and genomic data. Nature 562, 203–209.

36. Longchamps, R.J., Castellani, C.A., Yang, S.Y., Newcomb, C.E., Sumpter, J.A., Lane, J., Grove, M.L., Guallar, E., Pankratz, N., Taylor, K.D., et al. (2020). Evaluation of mitochondrial DNA copy number estimation techniques. PLOS ONE 15, e0228166.

37. MitoPipeline: Generating Mitochondrial copy number estimates from SNP array data in Genvisis.

38. Han, B., and Eskin, E. (2011). Random-Effects Model Aimed at Discovering Associations in Meta-Analysis of Genome-wide Association Studies. The American Journal of Human Genetics 88, 586–598.

39. Willer, C.J., Li, Y., and Abecasis, G.R. (2010). METAL: fast and efficient meta-analysis of genomewide association scans. Bioinformatics 26, 2190–2191.

40. Loh, P.-R., Tucker, G., Bulik-Sullivan, B.K., Vilhjálmsson, B.J., Finucane, H.K., Salem, R.M., Chasman, D.I., Ridker, P.M., Neale, B.M., Berger, B., et al. (2015). Efficient Bayesian mixed model analysis increases association power in large cohorts. Nat Genet 47, 284–290.

41. Speed, D. (2019). SumHer better estimates the SNP heritability of complex traits from summary statistics. Nature Genetics 51, 12.

42. Wang, G., Sarkar, A., Carbonetto, P., and Stephens, M. (2020). A simple new approach to variable selection in regression, with application to genetic fine mapping. Journal of the Royal Statistical Society: Series B (Statistical Methodology) 82, 1273–1300.

43. Wang, K., Li, M., and Hakonarson, H. (2010). ANNOVAR: functional annotation of genetic variants from high-throughput sequencing data. Nucleic Acids Research 38, e164–e164.

44. Sherry, S.T., Ward, M., and Sirotkin, K. (1999). dbSNP-database for single nucleotide polymorphisms and other classes of minor genetic variation. Genome Res 9, 677–679.

45. O’Leary, N.A., Wright, M.W., Brister, J.R., Ciufo, S., Haddad, D., McVeigh, R., Rajput, B., Robbertse, B., Smith-White, B., Ako-Adjei, D., et al. (2016). Reference sequence (RefSeq) database at NCBI: current status, taxonomic expansion, and functional annotation. Nucleic Acids Res 44, D733–745.

46. Giambartolomei, C., Vukcevic, D., Schadt, E.E., Franke, L., Hingorani, A.D., Wallace, C., and Plagnol, V. (2014). Bayesian Test for Colocalisation between Pairs of Genetic Association Studies Using Summary Statistics. PLOS Genetics 10, e1004383.

47. Võsa, U., Claringbould, A., Westra, H.-J., Bonder, M.J., Deelen, P., Zeng, B., Kirsten, H., Saha, A., Kreuzhuber, R., Kasela, S., et al. (2018). Unraveling the polygenic architecture of complex traits using blood eQTL metaanalysis. BioRxiv 447367.

48. Pers, T.H., Karjalainen, J.M., Chan, Y., Westra, H.-J., Wood, A.R., Yang, J., Lui, J.C., Vedantam, S., Gustafsson, S., Esko, T., et al. (2015). Biological interpretation of genome-wide association studies using predicted gene functions. Nature Communications 6, 5890.

49. Smyth, G., Hu, Y., Ritchie, M., Silver, J., Wettenhall, J., McCarthy, D., Wu, D., Shi, W., Phipson, B., Lun, A., et al. (2021). limma: Linear Models for Microarray Data (Bioconductor version: Release (3.12)).

50. Young, M.D., Wakefield, M.J., Smyth, G.K., and Oshlack, A. (2010). Gene ontology analysis for RNA-seq: accounting for selection bias. Genome Biology 11, R14.

51. Barbeira, A.N., Dickinson, S.P., Bonazzola, R., Zheng, J., Wheeler, H.E., Torres, J.M., Torstenson, E.S., Shah, K.P., Garcia, T., Edwards, T.L., et al. (2018). Exploring the phenotypic consequences of tissue specific gene expression variation inferred from GWAS summary statistics. Nat Commun 9, 1825.

52. Barbeira, A.N., Pividori, M., Zheng, J., Wheeler, H.E., Nicolae, D.L., and Im, H.K. (2019). Integrating predicted transcriptome from multiple tissues improves association detection. PLOS Genetics 15, e1007889.

53. Supek, F., Bošnjak, M., Škunca, N., and Šmuc, T. (2011). REVIGO summarizes and visualizes long lists of gene ontology terms. PLoS One 6, e21800.

54. Stiles, A.R., Simon, M.T., Stover, A., Eftekharian, S., Khanlou, N., Wang, H.L., Magaki, S., Lee, H., Partynski, K., Dorrani, N., et al. (2016). Mutations in TFAM, encoding mitochondrial transcription factor A, cause neonatal liver failure associated with mtDNA depletion. Mol Genet Metab 119, 91–99.

55. Kornblum, C., Nicholls, T.J., Haack, T.B., Schöler, S., Peeva, V., Danhauser, K., Hallmann, K., Zsurka, G., Rorbach, J., Iuso, A., et al. (2013). Loss-of-function mutations in MGME1 impair mtDNA replication and cause multisystemic mitochondrial disease. Nat Genet 45, 214–219.

56. El-Hattab, A.W., and Scaglia, F. (2013). Mitochondrial DNA depletion syndromes: review and updates of genetic basis, manifestations, and therapeutic options. Neurotherapeutics 10, 186–198.

57. Millard, L.A.C., Davies, N.M., Gaunt, T.R., Davey Smith, G., and Tilling, K. (2018). Software Application Profile: PHESANT: a tool for performing automated phenome scans in UK Biobank. Int J Epidemiol 47, 29–35.

58. Konopka, T. (2020). umap: Uniform Manifold Approximation and Projection.

59. Hahsler, M., Piekenbrock, M., Arya, S., and Mount, D. (2019). dbscan: Density Based Clustering of Applications with Noise (DBSCAN) and Related Algorithms.

60. Delaneau, O., Zagury, J.-F., Robinson, M.R., Marchini, J.L., and Dermitzakis, E.T. (2019). Accurate, scalable and integrative haplotype estimation. Nat Commun 10,.

61. Rubinacci, S., Delaneau, O., and Marchini, J. (2020). Genotype imputation using the Positional Burrows Wheeler Transform. PLOS Genetics 16, e1009049.

62. Auton, A., Abecasis, G.R., Altshuler, D.M., Durbin, R.M., Abecasis, G.R., Bentley, D.R., Chakravarti, A., Clark, A.G., Donnelly, P., Eichler, E.E., et al. (2015). A global reference for human genetic variation. Nature 526, 68–74.

63. Shen, L., Attimonelli, M., Bai, R., Lott, M.T., Wallace, D.C., Falk, M.J., and Gai, X. (2018). MSeqDR mvTool: A mitochondrial DNA Web and API resource for comprehensive variant annotation, universal nomenclature collation, and reference genome conversion. Hum Mutat 39, 806–810.

64. Gibellini, L., De Gaetano, A., Mandrioli, M., Van Tongeren, E., Bortolotti, C.A., Cossarizza, A., and Pinti, M. (2020). The biology of Lonp1: More than a mitochondrial protease. Int Rev Cell Mol Biol 354, 1–61.

65. GTEx Consortium (2013). The Genotype-Tissue Expression (GTEx) project. Nat Genet 45, 580–585.

66. Liu, T., Lu, B., Lee, I., Ondrovicová, G., Kutejová, E., and Suzuki, C.K. (2004). DNA and RNA binding by the mitochondrial lon protease is regulated by nucleotide and protein substrate. J Biol Chem 279, 13902–13910.

67. Sharma, M.R., Koc, E.C., Datta, P.P., Booth, T.M., Spremulli, L.L., and Agrawal, R.K. (2003). Structure of the mammalian mitochondrial ribosome reveals an expanded functional role for its component proteins. Cell 115, 97–108.

68. Shimizu, S., Narita, M., and Tsujimoto, Y. (1999). Bcl-2 family proteins regulate the release of apoptogenic cytochrome c by the mitochondrial channel VDAC. Nature 399, 483–487.

69. Vagdatli, E., Gounari, E., Lazaridou, E., Katsibourlia, E., Tsikopoulou, F., and Labrianou, I. (2010). Platelet distribution width: a simple, practical and specific marker of activation of coagulation. Hippokratia 14, 28–32.

70. PathCards : Factors involved in megakaryocyte development and platelet production Pathway and related pathways.

71. PathCards : Apoptosis and Autophagy Pathway and related pathways.

72. Yuan, Y., Ju, Y.S., Kim, Y., Li, J., Wang, Y., Yoon, C.J., Yang, Y., Martincorena, I., Creighton, C.J., Weinstein, J.N., et al. (2020). Comprehensive molecular characterization of mitochondrial genomes in human cancers. Nature Genetics 52, 342–352.

73. Mizumachi, T., Muskhelishvili, L., Naito, A., Furusawa, J., Fan, C.-Y., Siegel, E.R., Kadlubar, F.F., Kumar, U., and Higuchi, M. (2008). Increased Distributional Variance of Mitochondrial DNA Content Associated With Prostate Cancer Cells as Compared With Normal Prostate Cells. Prostate 68, 408–417.

74. Basel, D. (2020). Mitochondrial DNA Depletion Syndromes. Clin Perinatol 47, 123–141.

75. DeBarmore, B., Longchamps, R.J., Zhang, Y., Kalyani, R.R., Guallar, E., Arking, D.E., Selvin, E., and Young, J.H. (2020). Mitochondrial DNA copy number and diabetes: the Atherosclerosis Risk in Communities (ARIC) study. BMJ Open Diabetes Res Care 8,.

76. Yang, S.Y., Castellani, C.A., Longchamps, R.J., Pillalamarri, V.K., O’Rourke, B., Guallar, E., and Arking, D.E. (2021). Blood-derived mitochondrial DNA copy number is associated with gene expression across multiple tissues and is predictive for incident neurodegenerative disease. Genome Res.

77. Hannah-Shmouni, F., MacNeil, L., Brady, L., Nilsson, M.I., and Tarnopolsky, M. (2019). Expanding the Clinical Spectrum of LONP1-Related Mitochondrial Cytopathy. Front Neurol 10, 981.

78. Grainha, T.R.R., Jorge, P.A. da S., Pérez-Pérez, M., Pérez Rodríguez, G., Pereira, M.O.B.O., and Lourenço, A.M.G. (2018). Exploring anti-quorum sensing and anti-virulence based strategies to fight Candida albicans infections: an in silico approach. FEMS Yeast Res 18,.

79. Ali, A.T., Idaghdour, Y., and Hodgkinson, A. (2020). Analysis of mitochondrial m1A/G RNA modification reveals links to nuclear genetic variants and associated disease processes. Communications Biology 3, 1–11.

80. Schrepfer, E., and Scorrano, L. (2016). Mitofusins, from Mitochondria to Metabolism. Mol Cell 61, 683–694.

81. Ishihara, N., Eura, Y., and Mihara, K. (2004). Mitofusin 1 and 2 play distinct roles in mitochondrial fusion reactions via GTPase activity. J Cell Sci 117, 6535–6546.

82. Zhang, D., Xi, Y., Coccimiglio, M.L., Mennigen, J.A., Jonz, M.G., Ekker, M., and Trudeau, V.L. (2012). Functional prediction and physiological characterization of a novel short trans-membrane protein 1 as a subunit of mitochondrial respiratory complexes. Physiol Genomics 44, 1133–1140.

83. Cavdar Koc, E., Burkhart, W., Blackburn, K., Moseley, A., and Spremulli, L.L. (2001). The small subunit of the mammalian mitochondrial ribosome. Identification of the full complement of ribosomal proteins present. J Biol Chem 276, 19363–19374.

84. Márquez-Jurado, S., Díaz-Colunga, J., das Neves, R.P., Martinez-Lorente, A., Almazán, F., Guantes, R., and Iborra, F.J. (2018). Mitochondrial levels determine variability in cell death by modulating apoptotic gene expression. Nat Commun 9, 389.

85. Dölle, C., Flønes, I., Nido, G.S., Miletic, H., Osuagwu, N., Kristoffersen, S., Lilleng, P.K., Larsen, J.P., Tysnes, O.-B., Haugarvoll, K., et al. (2016). Defective mitochondrial DNA homeostasis in the substantia nigra in Parkinson disease. Nat Commun 7, 13548.

86. Chen, C., Turnbull, D.M., and Reeve, A.K. (2019). Mitochondrial Dysfunction in Parkinson’s Disease-Cause or Consequence? Biology (Basel) 8,.

87. Pinto, M., and Moraes, C.T. (2014). Mitochondrial genome changes and neurodegenerative diseases. Biochim Biophys Acta 1842, 1198–1207.

88. Rodríguez-Santiago, B., Casademont, J., and Nunes, V. (2001). Is mitochondrial DNA depletion involved in Alzheimer’s disease? Eur J Hum Genet 9, 279–285.

89. Chew, H., Solomon, V.A., and Fonteh, A.N. (2020). Involvement of Lipids in Alzheimer’s Disease Pathology and Potential Therapies. Front Physiol 11, 598.

90. Yin, J., Reiman, E.M., Beach, T.G., Serrano, G.E., Sabbagh, M.N., Nielsen, M., Caselli, R.J., and Shi, J. (2020). Effect of ApoE isoforms on mitochondria in Alzheimer disease. Neurology 94, e2404–e2411.

91. El Khoury, J.B., Moore, K.J., Means, T.K., Leung, J., Terada, K., Toft, M., Freeman, M.W., and Luster, A.D. (2003). CD36 mediates the innate host response to beta-amyloid. J Exp Med 197, 1657–1666.

92. Lämsä, R., Helisalmi, S., Herukka, S.-K., Tapiola, T., Pirttilä, T., Vepsäläinen, S., Hiltunen, M., and Soininen, H. (2008). Genetic study evaluating LDLR polymorphisms and Alzheimer’s disease. Neurobiol Aging 29, 848–855.

93. Tomioka, M., Toda, Y., Mañucat, N.B., Akatsu, H., Fukumoto, M., Kono, N., Arai, H., Kioka, N., and Ueda, K. (2017). Lysophosphatidylcholine export by human ABCA7. Biochim Biophys Acta Mol Cell Biol Lipids 1862, 658–665.

94. Nuytemans, K., Maldonado, L., Ali, A., John-Williams, K., Beecham, G.W., Martin, E., Scott, W.K., and Vance, J.M. (2016). Overlap between Parkinson disease and Alzheimer disease in ABCA7 functional variants. Neurol Genet 2, e44.

95. Crabos, M., Yamakado, T., Heizmann, C.W., Cerletti, N., Bühler, F.R., and Erne, P. (1991). The calcium binding protein tropomyosin in human platelets and cardiac tissue: elevation in hypertensive cardiac hypertrophy. Eur J Clin Invest 21, 472–478.

96. Kostan, J., Salzer, U., Orlova, A., Törö, I., Hodnik, V., Senju, Y., Zou, J., Schreiner, C., Steiner, J., Meriläinen, J., et al. (2014). Direct interaction of actin filaments with F-BAR protein pacsin2. EMBO Rep 15, 1154–1162.

97. Sever, S. (2002). Dynamin and endocytosis. Curr Opin Cell Biol 14, 463–467.

98. Cai, B., Giridharan, S.S.P., Zhang, J., Saxena, S., Bahl, K., Schmidt, J.A., Sorgen, P.L., Guo, W., Naslavsky, N., and Caplan, S. (2013). Differential roles of C-terminal Eps15 homology domain proteins as vesiculators and tubulators of recycling endosomes. J Biol Chem 288, 30172–30180.

99. Di Buduo, C.A., Moccia, F., Battiston, M., De Marco, L., Mazzucato, M., Moratti, R., Tanzi, F., and Balduini, A. (2014). The importance of calcium in the regulation of megakaryocyte function. Haematologica 99, 769–778.

100. De Botton, S., Sabri, S., Daugas, E., Zermati, Y., Guidotti, J.E., Hermine, O., Kroemer, G., Vainchenker, W., and Debili, N. (2002). Platelet formation is the consequence of caspase activation within megakaryocytes. Blood 100, 1310–1317.

101. Josefsson, E.C., James, C., Henley, K.J., Debrincat, M.A., Rogers, K.L., Dowling, M.R., White, M.J., Kruse, E.A., Lane, R.M., Ellis, S., et al. (2011). Megakaryocytes possess a functional intrinsic apoptosis pathway that must be restrained to survive and produce platelets. J Exp Med 208, 2017–2031.

102. Josefsson, E.C., Burnett, D.L., Lebois, M., Debrincat, M.A., White, M.J., Henley, K.J., Lane, R.M., Moujalled, D., Preston, S.P., O’Reilly, L.A., et al. (2014). Platelet production proceeds independently of the intrinsic and extrinsic apoptosis pathways. Nat Commun 5, 3455.

103. McArthur, K., Chappaz, S., and Kile, B.T. (2018). Apoptosis in megakaryocytes and platelets: the life and death of a lineage. Blood 131, 605–610.

104. Linsel-Nitschke, P., Götz, A., Erdmann, J., Braenne, I., Braund, P., Hengstenberg, C., Stark, K., Fischer, M., Schreiber, S., El Mokhtari, N.E., et al. (2008). Lifelong reduction of LDL-cholesterol related to a common variant in the LDL-receptor gene decreases the risk of coronary artery disease--a Mendelian Randomisation study. PLoS One 3, e2986.

105. Lawlor, D.A., Harbord, R.M., Sterne, J.A.C., Timpson, N., and Davey Smith, G. (2008). Mendelian randomization: using genes as instruments for making causal inferences in epidemiology. Stat Med 27, 1133–1163.

106. Sookoian, S., Rosselli, M.S., Gemma, C., Burgueño, A.L., Gianotti, T.F., Castaño, G.O., and Pirola, C.J. (2010). Epigenetic regulation of insulin resistance in nonalcoholic fatty liver disease: Impact of liver methylation of the peroxisome proliferator–activated receptor γ coactivator 1α promoter. Hepatology 52, 1992–2000.

107. Vyas, C.M., Ogata, S., Reynolds, C.F., Mischoulon, D., Chang, G., Cook, N.R., Manson, J.E., Crous-Bou, M., De Vivo, I., and Okereke, O.I. (2020). Lifestyle and behavioral factors and mitochondrial DNA copy number in a diverse cohort of mid-life and older adults. PLoS One 15, e0237235.

