## Supplemental Figures and Data for "Genetic analysis of mitochondrial DNA copy number and associated traits identifies loci implicated in nucleotide metabolism, platelet activation, and megakaryocyte proliferation, and reveals a causal association of mitochondrial function with mortality"

**Figure S1. Co-localization results for *TFAM*, before (left) and after (right) removing the SNP with the lowest combined mtDNA-CN GWAS and *TFAM* eQTL p-value.**

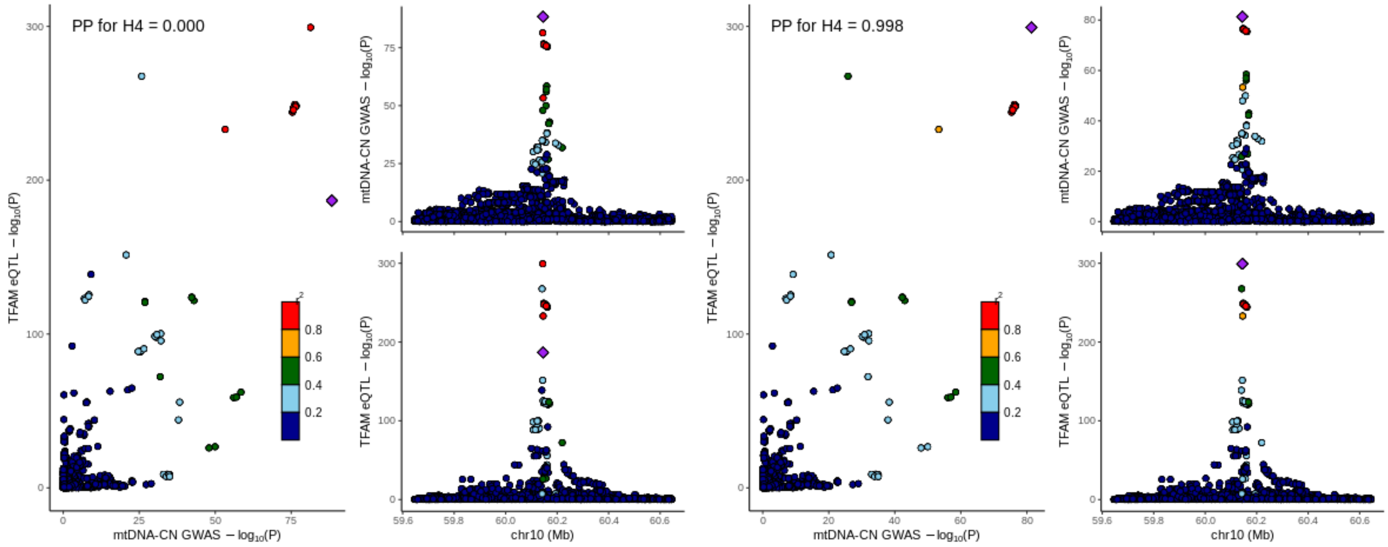


*TFAM* strongly colocalizes with GWAS hits after removing the SNP with the lowest combined mtDNA-GWAS and *TFAM* eQTL p-value. Colocalization was run with the R *coloc* package, and graphs were created using the *locusComparer* package.

**Figure S2. Flowchart describing gene prioritization process for each locus using results from ANNOVAR, SuSie, COLOC, and DEPICT.**

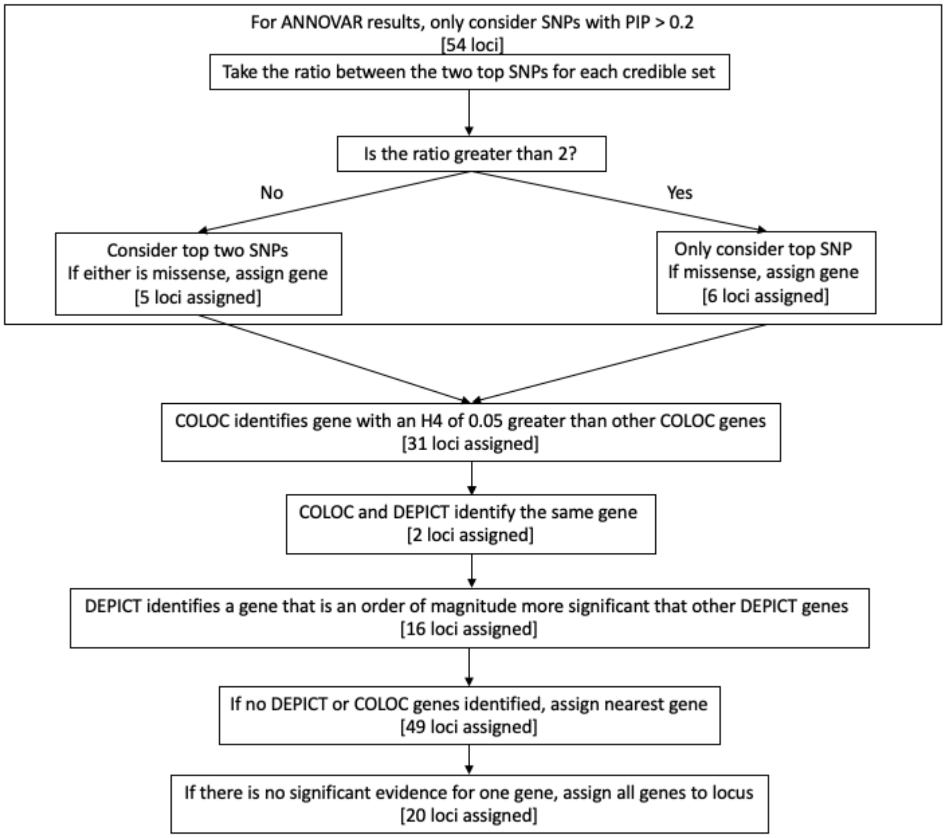


Flowchart depicting gene assignment pipeline. The number of loci assigned using each method is displayed in brackets.

**Figure S3. Primary autosomal loci identified in the 3 different complementary meta-analyses.**

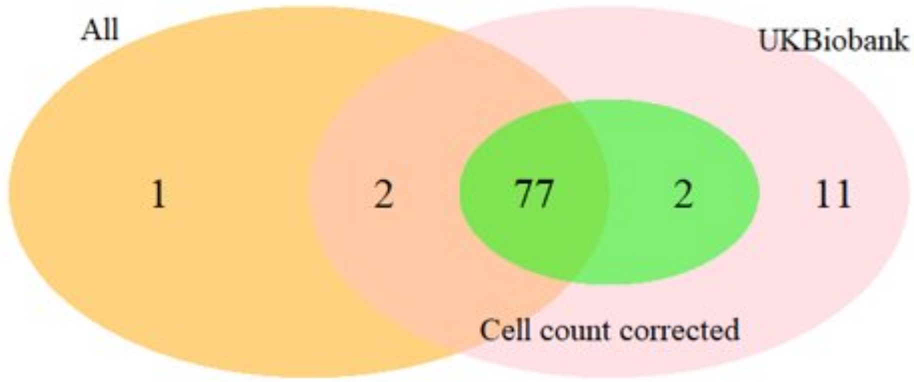


Primary autosomal loci identified in 1) All whites in the study (All), 2) All whites with available cell count information (cell count corrected), and 3) UKBiobank-only data (UKBiobank). The UKB-only GWAS identifies 92 of the 93 loci identified in all analyses, which is unsurprising as over 90% of samples come from the UKB cohort.

**Figure S4. QQ-plot showing expected and observed p-value distributions from autosomal UK Biobank GWAS summary statistics.**

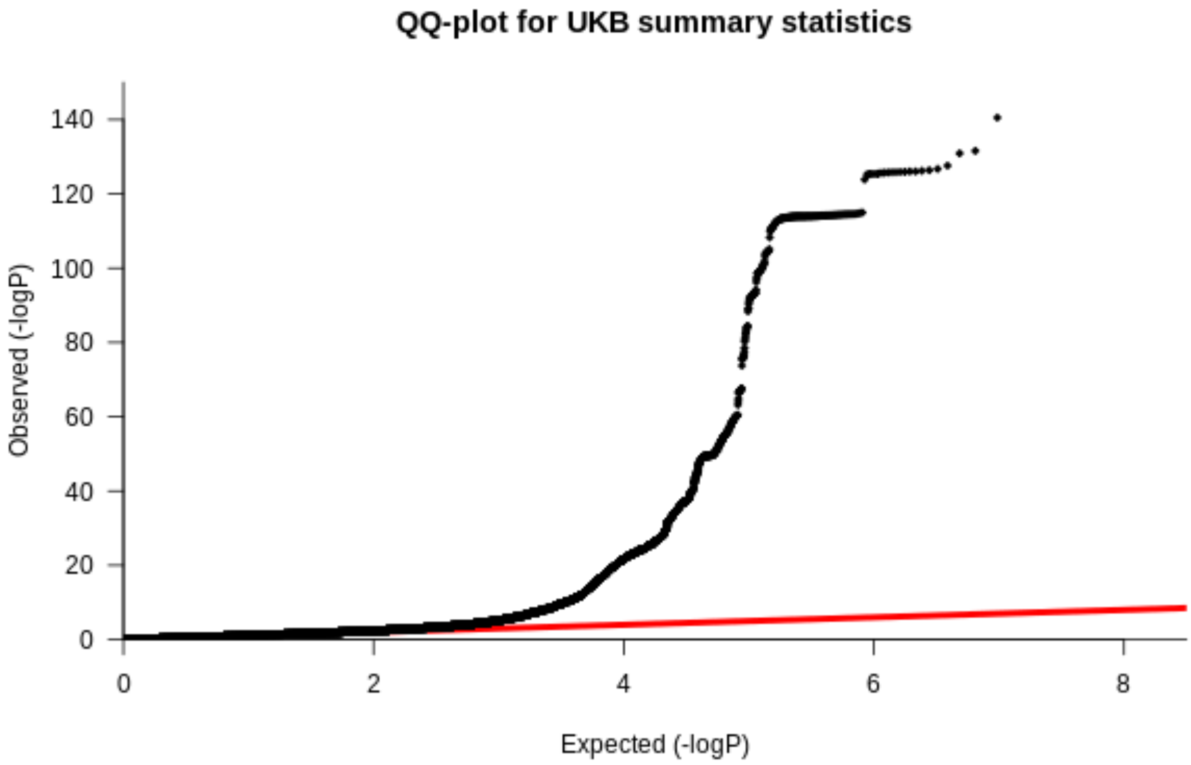

**Figure S5. SusieR fine-mapping results.**

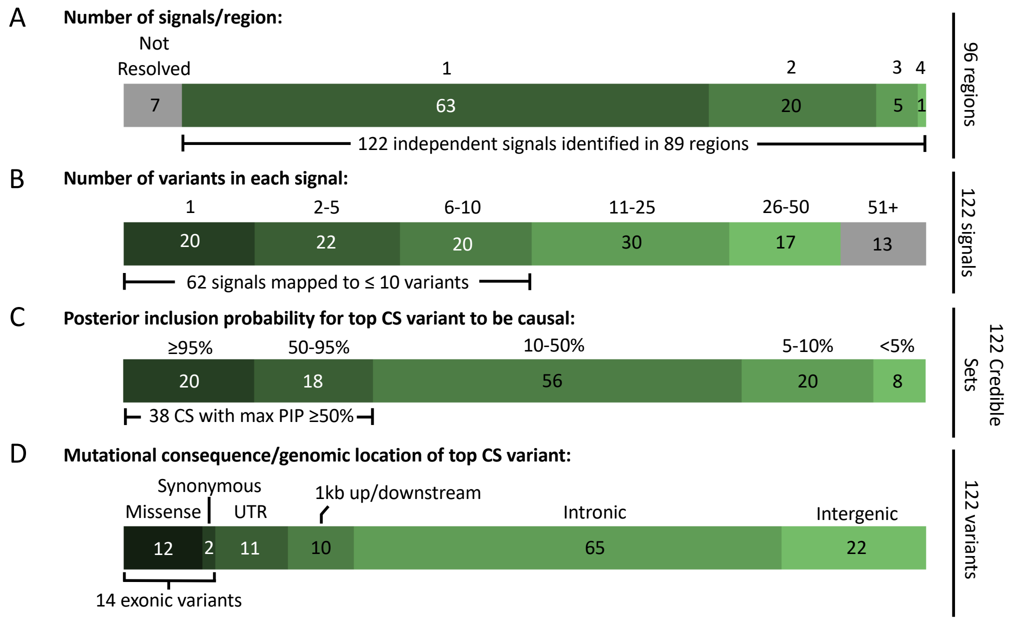


A. Out of 129 independent autosomal signals, susieR was able to resolve 122 credible sets in 89 regions. B. In each credible set, 62 credible sets were mapped to fewer than 10 variants. C. In 38 credible sets identified by susieR, the SNP with the highest PIP had a PIP greater than 50%. D. Of all the top SNPs for each credible set, 14 appeared in exonic regions, and 12 of those were missense mutations.

**Figure S6. ReviGO visualization of gene sets that are enriched for significantly associated MetaXcan genes.**

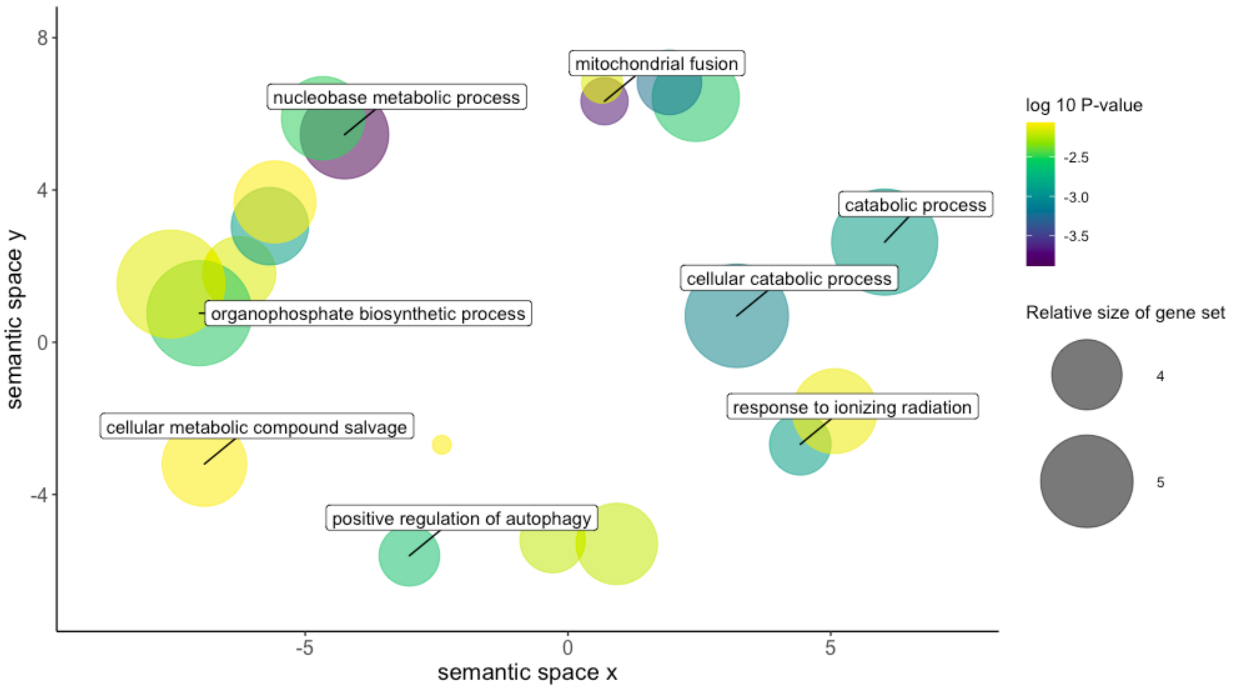


Gene sets that are significantly enriched in for genes with predicted gene expression that is associated with mtDNA-CN. Only terms with a p-value < 0.01 were included in this graph. X and Y axes represent semantic similarity between GO terms.

### Supplemental Methods

**UKB mtDNA-CN derivation.** We started with 49,997 Exome SPB CRAM files (version Jul 2018) downloaded from the UKB data repository, and used Samtools (ver1.9) to extract read summary statistics ('idxstats' command). A custom perl script was used to aggregate the summary statistics from each individual file into the following categories (see perl script and example stats file): 1) Total Reads (sum of columns 3 and 4, across all rows), 2) Mapped Reads (sum of column 3, across all rows), 3) Unmapped Reads (some of column 4 across all rows), 4) Autosomal Reads (sum of column 3, rows 1-22), 5) Chr X, 6) Chr Y, 7) Chr MT, 8) 'Random' Reads (sum of column 3, across rows 26-67), 9) 'Unknown' Reads (sum of column 3 across rows 68-194), 10) EBV Reads, 11) 'Decoy1' Reads (sum of column 3 across rows 196-582), 12) 'Decoy2' Reads (sum of column 3 across rows 583-2580). Linear regression models were used to adjust for total DNA and potential technical artifacts. Specifically, we used 10-fold cross validation for variable selection, using the 'leaps' R package (version 3.0), with an initial model with chrMT read count as the dependent variable, and 'Total', 'Mapped', 'unknown', 'random', 'decoy1' and 'decoy2' read counts as the independent variables. For each of the independent variables, we included a natural spline with df=4 to allow for non-linear effects. The independent variables 'Total', 'unknown', 'decoy1' and 'decoy2' read counts were selected. We then increased the natural spline df to 15, and then used backward selection to reduction model complexity, requiring P<0.005 to keep a term in the model. The final regression model residuals were generated with the following R (version 3.6.0) code:

```
WES.mtDNA=residuals(lm(chrMT ~ ns(Total,df=3) + ns(unknown,df=4) + ns(decoy1,df=7) + decoy2))
```

Mitochondrial SNP probe intensities were obtained from the "ukb_chrMT_l2r.txt" file downloaded from the UKBiobank, and samples were stratified by array type (UK BiLEVE, Axiom). To correct for potential artifacts and/or batch effects, we generated 250 genotyping principal components (PCs) using the 'rpca' command from the 'rsvd' package (version 1.0.3) from autosomal nuclear probes by randomly sampling 5% of probes from either even or odd chromosomes that were required to be present on both array types (n~19,500 probes). Note that we generated the two independent sets of PCs so that we could ensure that probe selection for PCA did not bias results. Prior to PCA, all probe intensities were rank transformed to reduce the impact of any outliers. For each array type, all mitochondrial SNP probes (UKBelieve, n=181; Axiom, n=244) along with the 250 PCs were regressed on the 'WES.mtDNA' metric derived as described above. Beta estimates from these analyses were then used to generate fitted values in the full UKBiobank dataset using the 'predict' function ('array.mtDNA').

Given the known impact of age, sex, and cell counts on mtDNA-CN, we first used visual inspection to identify outliers for cell counts:

Log(WBC) ≤1.25 or ≥3
Log(RBC) ≤1.4 or ≥2
Platelet ≤10 or ≥500
Log(Lymphocyte) ≤0.10 or ≥2
Log(Mono) ≥0.9
Log(Neutrophil) ≤0.75 or ≥2.75
Log(Eos) ≥0.75
Log(Baso) ≥0.45

We then excluded non-Whites, related individuals (used.in.pca.calculation=0), and cell count outliers and then adjusted for age, sex, and cell counts using a backwards regression, starting with a natural spline (df=4) for each covariate. The final model obtained was ("log_" indicates log-transformed variable):

mtDNA-CN = residuals(lm(array.mtDNA ~ ns(age,df=4) + sex + ns(log_WBC,df=4) + ns(log_RBC,df=4) + ns(Platelet,df=4) + ns(log_Lymph,df=4) + ns(log_Neutrophil,df=4) + log_Eos + log_Baso + log_NucRBC))

Beta estimates from these analyses were then used to generate fitted values in the full UKBiobank dataset using the 'predict' function.

For all analyses, mtDNA-CN was standardized by subtracting the mean and dividing by the standard deviation.

Table with variables from UKBB:

| UKB_ID | Name | Description |
|---|---|---|
| f.30000.0.0 | WBC | White blood cell (leukocyte) count |
| f.30010.0.0 | RBC | Red blood cell (erythrocyte) count |
| f.30080.0.0 | Platelet | Platelet count |
| f.30120.0.0 | Lymphocyte | Lymphocyte count |
| f.30130.0.0 | Mono | Monocyte count |
| f.30140.0.0 | Neutrophil | Neutrophil lcount |
| f.30150.0.0 | Eos | Eosinophill count |
| f.30160.0.0 | Baso | Basophill count |
| f.30170.0.0 | NucRBC | Nucleated red blood cell count |

**Cohort Descriptions**

*ALSPAC*

Pregnant women resident in Avon, UK with expected dates of delivery 1st April 1991 to 31st December 1992 were invited to take part in the study. The initial number of pregnancies enrolled is 14,541. Of these initial pregnancies, there was a total of 14,676 foetuses, resulting in 14,062 live births and 13,988 children who were alive at 1 year of age. When the oldest children were approximately 7 years of age, an attempt was made to bolster the initial sample with eligible cases who had failed to join the study originally. As a result, when considering variables collected from the age of seven onwards (and potentially abstracted from obstetric notes) there are data available for more than the 14,541 pregnancies mentioned above. The number of new pregnancies not in the initial sample (known as Phase I enrolment) that are currently represented on the built files and reflecting enrolment status at the age of 24 is 913 (456, 262 and 195 recruited during Phases II, III and IV respectively), resulting in an additional 913 children being enrolled. The phases of enrolment are described in more detail in the cohort profile paper and its update[1,2]. The total sample size for analyses using any data collected after the age of seven is therefore 15,454 pregnancies, resulting in 15,589 foetuses. Of these 14,901 were alive at 1 year of age. Please note that the study website contains details of all the data that is available through a fully searchable data dictionary and variable search tool" and reference the following webpage: http://www.bristol.ac.uk/alspac/researchers/our-data/

*ARIC*

The Atherosclerosis Risk in Communities study (ARIC) recruited 15,792 individuals between 1987 and 1989 aged 45 to 65 years from 4 US communities[3]. DNA for mtDNA-CN estimation was collected from different visits and was derived from buffy coat using the Gentra Puregene Blood Kit (Qiagen). Whole genome sequencing (WGS) data was generated at the Baylor College of Medicine Human Genome Sequencing Center using Nano or PCR-free DNA libraries on the Illumina HiSeq 2000. Genotyping for the Affymetrix Genome-Wide 6.0 Human SNP Array 6.0 was performed in accordance to the manufacturers protocol and genotypes were called using Birdseed (version 2).

With the availability of two different mtDNA-CN estimation platforms, and previous researching highlighting WGS outperforms the Affymetrix array in mtDNA-CN estimation[4], mtDNA-CN was called first from the 2,923 individuals with WGS data. Copy number was additionally estimated in 8,687 from the Affymetrix genotyping array for individuals without WGS data. WGS and Affymetrix mtDNA-CN batches were treated as separate cohorts due to differences in population distribution. Missing white blood cell count data (14.7%) was imputed to the mean.

*CHS*

The Cardiovascular Health Study (CHS) is a population-based cohort study of risk factors for coronary heart disease and stroke in adults ≥65 years conducted across four field centers[5]. The original predominantly White cohort of 5,201 persons was recruited in 1989-1990 from random samples of the Medicare eligibility lists; subsequently, an additional predominantly Black cohort of 687 persons were enrolled for a total sample of 5,888.

Blood samples were drawn from all participants at their baseline examination and DNA was subsequently extracted from available samples. Genotyping was performed at the General Clinical Research Center's Phenotyping/Genotyping Laboratory at Cedars-Sinai among CHS participants who consented to genetic testing and had DNA available using the Illumina 370CNV BeadChip system (for White participants, in 2007) or the Illumina HumanOmni1-Quad_v1 BeadChip system (for Black participants, in 2010).

*MESA*

The MESA study recruited 6,814 individuals from 6 US communities free of prevalent clinical CVD across 4 ethnicities. Age range at baseline was 45 to 84 and the baseline exam occurred between 2000 and 2002. Affymetrix Genome-Wide Human SNP Array 6.0 genotype data was available for 8,227 unique individuals within the MESA cohort. DNA derived from MESA Family, a subset of MESA, originated from cell lines and was excluded resulting in a final sample size of 5,916. DNA for mtDNA-CN analyses was isolated from exam 1 peripheral leukocytes using the Gentra Puregene Blood Kit.

*ROS/MAP*

The Rush Religious Orders Study (ROS), started in 1994, enrolled Catholic priests, nuns, and brothers, from about 45 groups in 14 states. Since January 1994 to the time this dataset was frozen, 1321 participants completed their baseline evaluation, of whom 1259 were non-Hispanic white. The follow-up rate of survivors exceeds 90%. Participants were free of known dementia at enrollment, agreed to annual clinical evaluations, and signed both an informed consent and an Anatomic Gift Act donating their brains at time of death. They also signed a repository consent allowing their data to be shared. ROS was approved by an Institutional Review Board of Rush University Medical Center. A more detailed description of ROS has been published previously[6]. Participation included a neuropsychological test battery. DNA was extracted from whole blood or brain. Genotyping was performed at the Broad Institute's Center for Genotyping and the Translational Genomics Research Institute.

The Rush Memory and AP (MAP), started in 1997, enrolled older men and women from retirement communities, subsidized housing, and individual home in the greater Chicago area without known dementia at baseline. Since October 1997 to the time this dataset was frozen, 1815 participants completed their baseline evaluation, of whom 1701 were non-Hispanic white people. The follow-up rate of survivors exceeds 90%. Participants agreed to annual clinical evaluations, and signed both an informed consent and an Anatomic Gift Act donating their brains at time of death. They also signed a repository consent allowing their data to be shared. ROS was approved by an Institutional Review Board of Rush University Medical Center. A more detailed description of the MAP has been published previously[7]. Participation included a neuropsychological test battery. DNA was extracted from whole blood or brain. Genotyping was performed at the Broad Institute's Center for Genotyping and the Translational Genomics Research Institute.

*SHIP*
The Study of Health in Pomerania (SHIP) is a population-based project in West Pomerania, the north-east area of Germany[8,9]. A sample from the population aged 20 to 79 years was drawn from population registries. First, the three cities of the region (with 17,076 to 65,977 inhabitants) and the 12 towns (with 1,516 to 3,044 inhabitants) were selected, and then 17 out of 97 smaller towns (with less than 1,500 inhabitants), were drawn at random. Second, from each of the selected communities, subjects were drawn at random, proportional to the population size of each community and stratified by age and gender. Only individuals with German citizenship and main residency in the study area were included. Finally, 7,008 subjects were sampled, with 292 persons of each gender in each of the twelve five-year age strata. In order to minimize drop-outs by migration or death, subjects were selected in two waves. The net sample (without migrated or deceased persons) comprised 6,267 eligible subjects. Selected persons received a maximum of three written invitations. In case of non-response, letters were followed by a phone call or by home visits if contact by phone was not possible. The SHIP population finally comprised 4,308 participants (corresponding to a final response of 68.8%).

*UK Biobank*
The UK Biobank is a prospective cohort of over 500,000 individuals originating from 22 centers within the United Kingdom aged between 40 and 69.[10] Biological samples and physical measurements were collected and participants answered extensive questionnaires on health. Systolic and diastolic blood pressures are the average of two measurements taken at baseline. Genotyping was performed with the Affymetrix Axiom array.

**Cohort Specific QC and GWAS**
*ALSPAC*
Genotypes derived from the Illumina Human660W-Quad array. Individuals were dropped if they had > 5% missing, outliers for autosomal heterozygosity, indeterminate X chromosome, cryptic relatedness > 0.125, ID mismatch or non-European ancestry. SNPs were dropped if they had missingness > 5%, minor allele frequency < 0.01, or an imputation INFO score < 0.8. Haplotypes were estimated using ShapeIT (v2.r644), and relatedness was utilized during phasing. Phased haplotypes were imputed to the

Haplotype Reference Consortium (HRC). The HRC panel itself was phased using ShapeIT (v2), with imputation performed using IMPUTE (v3).

GWAS was additionally adjusted for 2 genotyping principal components.

*ARIC*
Genotypes derived from the Affymetrix Genome-Wide Human SNP Array 6.0. Individuals were dropped if they refused DNA testing, > 5% missingness, or were identified as genetic outliers. Monomorphic SNPs, and SNPs with > 5% missingness were dropped. Whites were imputed to the Haplotype Reference Consortium while Blacks were imputed to 1000G (March 2012).

GWAS was additionally adjusted for 10 genotyping principal components.

*CHS*
Genotypes for Whites were derived from the Illumina 370CNV chip, ITMAT-Broad-CARe Illumina iSELECT chip. Individuals were excluded from the sample due to the presence of baseline CHD, CHF, peripheral vascular disease, valvular heart disease, stroke or TIA, or lack of available DNA. After genotyping, individuals were excluded if they had a call rate <=95% or if their genotype was discordant with known sex or prior genotyping. SNPs were excluded if they had a call rate <97%, HWE P < 10-5, >2 duplicate errors or Mendelian inconsistencies, or heterozygote frequency = 0.

Genotypes for Black individuals were derived from the Illumina HumanOmni1-Quad_v1 BeadChip system. Individuals were excluded from the sample due to lack of available DNA. After genotyping, individuals were excluded if they had a call rate <=95% or if their genotype was discordant with known sex or prior genotyping. SNPs were excluded if they had a call rate <97%, HWE P < 10-5, >1 duplicate error or Mendelian inconsistency, or heterozygote frequency = 0.

Blacks-only GWAS was additionally adjusted for 5 genotyping principal components.

*FHS*
Genotyping derived from the Affymetrix 500K mapping array plus Affymetrix 50K supplemental array. Genotyped SNPs removed based on: HWE p-value of less than 0.000001, call rate of less than 96.9%, MAF of less than 0.01, no mapping correctly from Build 36 to Build 37 locations, missing a physical location, Mendelian errors greater than 1000, not being on chromosomes 1-22 or X, duplicates. Imputed to 1000G (March 2012).

Due to family structure, a linear mixed effects model with fixed polygenic variance was used to account for family relationship during the GWAS.

*MESA*
Genotypes derived from the Affymetrix Genome-Wide Human SNP Array 6.0. Individuals with < 95% genotyping success rate were excluded while SNPs with < 90% genotyping success rate were excluded. Imputation was performed to 1000G (March 2012).

GWAS was additionally adjusted for 4 genotyping principal components.

*ROS/MAP*

Genotypes derived from the Affymetrix Genome-Wide Human SNP Array 6.0. Sample-level quality control assessment included exclusion of samples with genotype success rate <95%, discordance between inferred and reported gender, and excess inter/intraheterozygosity. SNP-level quality control assessment included exclusion of SNPs with Hardy-Weighberg equilibrium ($p<0.001$), MAF < 0.01, genotype call rate < 0.95, misshap test < $1x10^{-9}$. Subsequently, EIGENSTRAT was used to identify and remove population outliers using default parameters

*SHIP*

Genotypes derived from the Affymetrix Genome-Wide Human SNP Array 6.0. Arrays with a call rate below 94%, duplicate samples as identified by estimated IBD as well as individuals with reported and genotyped gender mismatch were excluded. The final sample call rate was 99.51%.

GWAS was additionally adjusted for 10 PCs to account for population substructure.

*UK Biobank*

Whole exome sequencing, sequencing alignment and variant identification was performed as previously described[11]. Genotypes were available through the Affymetrix UK Biobank Axiom array and the Affymetrix UK BiLEVE Axiom array. Quality control, phasing and imputation information have been described previously (http://biobank.ctsu.ox.ac.uk/crystal/refer.cgi?id=155580 , (http://biobank.ctsu.ox.ac.uk/crystal/refer.cgi?id=157020).

Due to its recruitment method and extremely large sample size, as much as 30% of the UK Biobank dataset would need to be filtered due to typical relatedness or genetic ancestry filters[12]. To account for this, GWAS was performed using BOLT-LMM (v2.3.2) with the addition of a kinship matrix to control for related individuals[13]. GWAS was performed on all self-identified white individuals, excluding individuals who were cell count outliers as described above.

**Ethics Statements**

*ALSPAC*

Ethical approval for the study was obtained from the ALSPAC Ethics and Law Committee and the Local Research Ethics Committees. Informed consent for the use of data collected via questionnaires and clinics was obtained from participants following the recommendations of the ALSPAC Ethics and Law Committee at the time

*ARIC*

Institutional Review Board approvals were obtained by the coordinating center and each ARIC study center. The research was conducted in accordance with the principles described in the Helsinki Declaration. All subjects in the ARIC study gave informed consent.

*CHS*

CHS was approved by institutional review committees at each field center and individuals in the present analysis had available DNA and gave informed consent including consent to use of genetic information for the study of cardiovascular disease.

*FHS*

The Boston University Medical Campus Institutional Review Board approved the FHS genome-wide genotyping.

*MESA*
MESA All MESA participants provided written and informed consent to participate in genetic studies. All study sites received approval to conduct this research from local Institutional Review Boards.

*ROS/MAP*
All participants provided written informed consent and approval was obtained from an institutional review board. Participants also signed a repository consent to allow their data to be shared.

*SHIP*
The study has been conducted according to the recommendations of the Declaration of Helsinki. The study protocol of SHIP was approved by the medical ethics committee of the University of Greifswald. Written informed consent was obtained from each of the study participants.

*UK Biobank*
Research was approved by UK Biobank to ensure consistent with participant's consent and framework of data access.

**Supplemental References**


1. Boyd A, Golding J, Macleod J, et al. Cohort Profile: the 'children of the 90s'--the index offspring of the Avon Longitudinal Study of Parents and Children. *Int J Epidemiol*. 2013;42(1):111-127. doi:10.1093/ije/dys064

2. Fraser A, Macdonald-Wallis C, Tilling K, et al. Cohort Profile: the Avon Longitudinal Study of Parents and Children: ALSPAC mothers cohort. *Int J Epidemiol*. 2013;42(1):97-110. doi:10.1093/ije/dys066

3. The Atherosclerosis Risk in Communities (ARIC) Study: design and objectives. The ARIC investigators. *Am J Epidemiol*. 1989;129(4):687-702.

4. Longchamps RJ, Castellani CA, Yang SY, et al. Evaluation of mitochondrial DNA copy number estimation techniques. *PLOS ONE*. 2020;15(1):e0228166. doi:10.1371/journal.pone.0228166

5. Fried LP, Borhani NO, Enright P, et al. The Cardiovascular Health Study: design and rationale. *Ann Epidemiol*. 1991;1(3):263-276.

6. Bennett DA, Schneider JA, Arvanitakis Z, Wilson RS. OVERVIEW AND FINDINGS FROM THE RELIGIOUS ORDERS STUDY. *Curr Alzheimer Res*. 2012;9(6):628-645.

7. Bennett DA, Schneider JA, Buchman AS, Barnes LL, Boyle PA, Wilson RS. Overview and Findings from the Rush Memory and Aging Project. *Curr Alzheimer Res*. 2012;9(6):646-663.

8. John U, Greiner B, Hensel E, et al. Study of Health In Pomerania (SHIP): a health examination survey in an east German region: objectives and design. *Soz Praventivmed*. 2001;46(3):186-194.

9. Völzke H, Alte D, Schmidt CO, et al. Cohort profile: the study of health in Pomerania. *Int J Epidemiol*. 2011;40(2):294-307. doi:10.1093/ije/dyp394

10. Bycroft C, Freeman C, Petkova D, et al. The UK Biobank resource with deep phenotyping and genomic data. *Nature*. 2018;562(7726):203-209. doi:10.1038/s41586-018-0579-z

11. Whole exome sequencing and characterization of coding variation in 49,960 individuals in the UK Biobank | bioRxiv. Accessed August 14, 2019. https://www.biorxiv.org/content/10.1101/572347v1.supplementary-material

12. Loh P-R, Kichaev G, Gazal S, Schoech AP, Price AL. Mixed model association for biobank-scale data sets. *Nat Genet*. 2018;50(7):906-908. doi:10.1038/s41588-018-0144-6

13. Loh P-R, Tucker G, Bulik-Sullivan BK, et al. Efficient Bayesian mixed model analysis increases association power in large cohorts. *Nat Genet*. 2015;47(3):284-290. doi:10.1038/ng.3190
